# Eye movements reveal age differences in how arousal modulates saliency priority but not attention processing speed

**DOI:** 10.1101/2024.05.06.592619

**Authors:** Andy Jeesu Kim, Kristine Nguyen, Mara Mather

## Abstract

The arousal-biased competition theory posits that inducing arousal increases attentional priority of salient stimuli while reducing priority of non-pertinent stimuli. However, unlike in young adults, older adults rarely exhibit shifts in priority under increased arousal, and prior studies have proposed different neural mechanisms to explain how arousal differentially modulates selective attention in older adults. Therefore, we investigated how the threat of unpredictable shock differentially modulates attentional control mechanisms in young and older adults by observing eye movements. Participants completed two oculomotor search tasks in which the salient distractor was typically captured by attention (singleton search) or proactively suppressed (feature search). We found that arousal did not modulate attentional priority for any stimulus among older adults nor affect the speed of attention processing in either age group. Furthermore, we observed that arousal modulated pupil sizes and found a correlation between evoked pupil responses and oculomotor function. Our findings suggest age differences in how the locus coeruleus-noradrenaline system interacts with neural networks of attention and oculomotor function.

**Highlights:**

– Increased arousal modulates attention saliency priority but not processing speed
– Older adults do not exhibit shifts in stimulus priority under elevated arousal
– Proactive suppression of salient stimuli persists even during increased arousal
– Threat of unpredictable shock increases pupil sizes and decreases evoked responses
– Eye movements may be able to help assess locus coeruleus function

## Introduction

The brain’s visual and attention systems work in tandem to ensure the perception of pertinent information. The biased competition theory of attention posits that objects in the visual field compete for limited cognitive resources (Desimone & Duncan, 1995) through the activation and coordination of multiple neural networks of attention (Corbetta et al., 2008; Corbetta & Shulman, 2002). Biased competition thus favors behaviorally relevant stimuli such as objects that are salient, contain features relevant to task goals, and stimuli that have gained priority because of prior exposure, learning, and selection (Anderson et al., 2021). However, the attentional priority of stimuli and objects is not always constant in real-world scenarios, and changes in brain state (e.g., elevated arousal) can cause shifts in visual priority maps (Zelinsky & Bisley, 2015).

The locus coeruleus-noradrenaline (LC-NE) system mediates global arousal in the brain and influences cognitive processes, including attentional control, through its widespread efferent projections (Berridge, 2008; Berridge & Waterhouse, 2003; Maness et al., 2022). The Arousal-Biased Competition theory posits that inducing arousal elevates or impairs the likelihood of an object’s neural representation based on its given priority, highlighting a “winner-take-more” and “loser-take-less” selection effect in perception and memory (Mather & Sutherland, 2011).

Moreover, the Glutamate Amplifies Noradrenergic Effect (GANE) model poses that the biological basis of increased priority is because of local noradrenergic “hot spots” in which the co-release of excitatory neurotransmitter glutamate and norepinephrine signal elevated attentional priority (Mather et al., 2016b, 2016a). Importantly, this model also poses that widespread noradrenaline release outside of these local hot spots produces an inhibitory effect, addressing findings of arousal-induced suppression that are not accounted for in other prominent emotion-cognition theories (Mather et al., 2016b). Indeed, a recent systematic review found that participants allocate higher attentional priority to arousal-inducing stimuli in visual attention tasks and often experience improved performance when arousal is induced immediately before the task (Zsidó, 2023). These conclusions were consistent across multiple types of stimuli (e.g., emotional faces, affective images, auditory) and across visual search, attentional blink, and spatial cuing tasks.

While arousal leads to short-term changes in brain state, aging leads to sustained changes in mechanisms of selective attention. Compared with young adults, older adults commonly exhibit increases in distractibility by salient stimuli that are thought to be a byproduct of declines in mechanisms of inhibitory processing (Hasher & Zacks, 1988; Kubo-Kawai & Kawai, 2010; Lustig et al., 2007; West & Alain, 2000; Zeef et al., 1996). These behavioral changes are primarily associated with reduced functional connectivity and activation in the frontoparietal attention networks (Campbell et al., 2012; Geerligs et al., 2015; Grady et al., 2016; Kennedy & Mather, 2019; Madden, 2007; Nashiro et al., 2017) with some tasks increasing prefrontal cortex activity more in older than in younger adults, potentially as the older brain compensates for reduced function (Zanto & Gazzaley, 2017). However, studies using the Attention Network Test have identified age equivalence in mechanisms of orienting and conflict resolution but found that older adults exhibit impaired executive function and alerting compared with young adults (Gamboz et al., 2010; Mahoney et al., 2010; see also McDonough et al., 2019, for a review). Furthermore, a recent study has showed that aging improves the efficiency of the orienting and executive networks of attention (Veríssimo et al., 2022). These findings show that tasks requiring inhibition of salient stimuli and goal-directed spatial cuing result in impaired attentional control in older adults, but their ability may be preserved in orienting or flanker-type tasks administered in the Attention Network test. To explain these task-specific findings, Kim, Senior, et al. (2024) investigated how aging modulated multiple mechanisms of attentional control by examining the timing and direction of eye movements in two oculomotor search tasks. The authors found that older adults maintain preserved mechanisms of orienting during feature search and exhibit impaired goal-directed attentional control with the alerting cue as seen in the Attention Network Test, in addition to being significantly distracted by the salient distractor as in the aging and inhibition literature. Furthermore, older adults displayed age equivalence in mechanisms of selection history as previously observed (Howard Jr. et al., 2004; Lega et al., 2023; Lyon et al., 2014; Smyth & Shanks, 2011) and interestingly, could inhibit the salient distractor when the task design required proactive suppression. Although significant advances have been made to understand how aging affects mechanisms of attentional control, the neurobiological bases of why specific networks of attention are affected and why others remain unaltered is yet unknown.

One reason specific networks of attentional control may become impaired or are unaffected in aging may be because aging leads to functional impairments of the locus coeruleus-noradrenaline (LC-NE) system. The LC-NE system regulates cognitive processes, including attention (Aston-Jones et al., 1999, 2000; Bouret & Sara, 2004; Ghosh & Maunsell, 2022; Maness et al., 2022; Unsworth & Robison, 2017) and a foundation for healthy aging is an intact LC-NE system (Clewett et al., 2016; Mather & Harley, 2016; Wilson et al., 2013).

However, neuroimaging studies have revealed that older adults exhibit reduced locus coeruleus magnetic resonance imaging (MRI) contrast compared with young adults, showing changes in neuronal density (Cassidy et al., 2022; Clewett et al., 2016; Dahl et al., 2019, 2022, 2023; Hämmerer et al., 2018; K. Y. Liu et al., 2020). In older adults, reduced LC-MRI contrast is associated with decreased cognitive function and increased risk for the progression of neurodegenerative diseases (Elman et al., 2021; Hämmerer et al., 2018). Functional connectivity studies also reveal that older adults have altered connectivity between the LC and other brain regions compared with young adults (Jacobs et al., 2015; T.-H. Lee et al., 2018, 2020; Takahashi et al., 2015; Zhang et al., 2016). Collectively, these findings portray a dysfunctional LC-NE system in the aging human brain that is negatively affecting cognitive functions, including attentional selection. Researchers have hypothesized that noradrenergic hyperactivity, or elevated tonic (baseline) noradrenergic firing rates, characterizes this LC-NE systemic dysfunction in older adults compared with those of young adults (Elman et al., 2017; Gannon & Wang, 2019; Mather, 2020; Weinshenker, 2018). Based on the Yerkes-Dodson relationship between tonic and phasic LC activity (Aston-Jones & Cohen, 2005), tonic noradrenergic hyperactivity in older adults would produce decreased phasic noradrenaline release and sub-optimal performance.

In young adults with “unaltered” tonic noradrenergic release, increased arousal improves task performance, speed, and perceptual contrast (Laretzaki et al., 2010; T.-H. Lee et al., 2012; Padmala & Pessoa, 2008; Phelps et al., 2006; Schupp et al., 2007; Vuilleumier et al., 2001), in addition to memory recall (Clewett et al., 2018; Jia et al., 2020; Sakaki et al., 2019). The aforementioned GANE model proposes that the observed benefits in task performance in young adults are because of arousal-induced local noradrenaline release that amplifies attentional priority of pertinent stimuli (Mather et al., 2016b). Furthermore, researchers have shown that inducing arousal through the threat of shock reduces attentional priority of reward-related stimuli during visual search despite increased neural activity (Kim & Anderson, 2020b, 2020a), providing a prime example of how elevated arousal leads to a “loser-take-less” effect for stimuli with learned associations (e.g., reward) that are not physically salient and thus lower priority in the lens of the attention systems when threatened (Mather & Sutherland, 2011). However, if we hypothesize that older adults exhibit noradrenergic hyperactivity and the LC-NE system mediates attentional control, does inducing arousal in older adults modulate mechanisms of attentional control differently compared to young adults? Most studies in the literature reveal that arousal increases attention processing to salient or valenced stimuli in young adults but not in older adults (Allen et al., 2017; Durbin et al., 2018; Gallant et al., 2022; T.-H. Lee et al., 2018; Svärd et al., 2014; but see Sutherland & Mather, 2015), supporting the noradrenergic dysfunction hypothesis in aging. However, the neural mechanisms of how arousal-biased competition does not elevate priority of salient stimuli in older adults as much as in younger adults is not yet clear. Lee et al. (2018) observe that arousal increases neural activity of both salient and non-salient stimuli in older adults, thus leading to a lack of sensitivity for high salient stimuli. On the other hand, Gallant et al. (2022) observe that arousal increases sensitivity for high salience stimuli only in young adults but not in older adults, demonstrating age differences in selectivity based on the priority of a given stimulus.

To investigate age differences in mechanisms of arousal-biased competition, we used an oculomotor visual search task that can measure multiple mechanisms of attentional control including specific changes in attentional priority of salient and non-salient stimuli by observing eye movements (Kim, Senior, et al., 2024). Initiation of eye movements in visual attention and perception involves a neural network, including the frontal eye fields, visual cortical region V4, superior colliculus, and the mediodorsal thalamus (Armstrong et al., 2006; Bruce & Goldberg, 1985; Sommer & Wurtz, 2002, 2006). Efferent axons from the locus coeruleus end on all of these brain regions (Kalwani et al., 2014; Li et al., 2018; Loughlin et al., 1986; McBurney-Lin et al., 2019; Pickel et al., 1974; Schwarz et al., 2015) and the LC-NE system has been shown to modulate eye movements through this brain network (Grefkes et al., 2010; Kalwani et al., 2014; M. Lee et al., 2020; Sawaguchi & Kikuchi, 1998). We used the threat of unpredictable shock paradigm to induce arousal in both young and older adults, administering an electric shock at random times to emulate states of elevated tonic noradrenergic activity (Schmitz & Grillon, 2012). Prior studies have shown that the threat of unpredictable shock increases attentional priority of salient stimuli, decreases attentional priority of reward-associated stimuli, and also improves goal-directed attentional control in young adults (Kim et al., 2021; Kim & Anderson, 2020b, 2020a).

In the current study, we hypothesized that older adults would not show changes in attentional priority for all stimuli under threat of unpredictable shock, unlike young adults. If arousal increases first saccade destinations to only the salient distractor and not to non-salient stimuli in young adults but not in older adults, these results will support the findings of Gallant et al. (2022) and show specific shifts in priority based on the stimulus. However, if arousal increases first saccade destinations to both the salient distractor and non-salient stimuli in older adults, these findings will support the conclusions of Lee et al. (2018) and show general increases in bottom-up processing that is non-specific to salient stimuli. Besides investigating changes in attentional priority of three types of stimuli (first saccade destinations to either the target, non-target, salient distractor), we also examined whether increased arousal would differentially modulate the timing and speed of attentional processes and overall performance between young and older adults by investigating saccadic latencies (time to initiate the first saccade), fixation times (time to orient to the target), and dwell times (time to disengage after rejection of a non-target stimulus). Last, we explored these effects in two unique oculomotor search tasks: the first task required participants to fixate on the singleton shape (singleton-search) in the search array requiring reactive disengagement of the distractor while the second task required participants to fixation on a specific shape (feature-search) that leads to proactive suppression of a salient distractor (Kim, Senior, et al., 2024). We were interested in whether arousal-induced changes in attentional priority of the salient distractor would persist (at least in young adults) even if participants proactively suppressed the distractor, or whether increases in attentional priority would disrupt mechanisms of inhibition.

## Methods

### Participants and Sample Size Calculations

In this experiment, our first research goal was to investigate age differences in mechanisms of attentional control. The findings for this aim were reported in Kim, Senior, et al. (2024). Our second research goal was to investigate whether arousal would differentially modulate attentional priority of salient stimuli in young and older adults and the findings for this aim are reported in this manuscript. When preparing our first manuscript, eight participants did not complete the shock block and we observed a marginal interaction effect of arousal on age. Given that our original power analysis was to investigate age differences for our first research goal, we considered the possibility that we were underpowered to observe arousal by age interaction effects. Therefore, we conducted a second power analysis for our second research question.

To our knowledge, the unpredictable threat of shock paradigm has never been used in studies measuring attentional control within older adults. Similarly, Lee et al. (2018) induced arousal with an auditory tone conditioned with electric shock and observed a significant age x saliency type x arousal interaction effect. When examining the effect of arousal for each group, both young and older adults showed either a significant main effect of arousal or an interaction between arousal and saliency, with at least an effect size of η*_p_^2^* = .178. Using this large effect size (*f* = 0.4), we conducted a power analysis (G*Power 3.1) and identified that to observe a within-between interaction in a MANOVA analysis at α = 0.05 and power (1-β) = 0.9 a sample size of at least thirty-four young and older adults was required.

Thus, to be adequately powered to observe arousal-induced changes in attentional control, we newly recruited twenty young adults and sixteen older adults to add to our dataset from the first phase of data collection (eight participants in Kim, Senior, et al. (2024) did not complete the shock block and thus their data is not reported here). After the second phase of data collection, we recruited a total of thirty-four young adults (YA: female = 23, male = 10, prefer not to say = 1) aged 18-30 inclusive (mean = 21.2, SD = 2.9) and a total of thirty-four older adults (OA: female = 26, male = 8) aged 55-80 inclusive (mean = 69.0, SD = 7.2) to complete the singleton-search task. In addition, we recruited a sample of thirty-six young adults (YA: female = 25, male = 12) aged 18-23 inclusive (mean = 20.1, SD = 1.5) and thirty-six older adults (OA: female = 24, male = 11) aged 51-80 inclusive (mean = 68.2, SD = 8.0) to complete the feature-search task. Young adults were recruited from the University of Southern California (USC) SONA subject pool for course credit and older adults were recruited from the local Los Angeles communities for monetary compensation. Older adults were screened for cognitive dysfunction using the TELE screening protocol and individuals who scored below the cutoff threshold of 16 were excluded from the study (Gatz et al., 1995). All study protocols were approved by the USC Institutional Review Board and all participants provided written, informed consent prior to participation.

Although our results did not change after the first and second phases of data collection, we report our data collection procedure here for transparency. Despite some of our primary statistical analyses revealing marginal age by arousal interaction effects, we stopped data collection and report our findings based on our power analysis and to prevent the likelihood of false positives. We conducted Bayesian analyses when appropriate to explore evidence in support of the null hypothesis. Given that some of the non-arousal data (blocks without the threat of shock) has previously been reported in Kim, Senior, et al. (2024), the age-related findings in both manuscripts should not be interpreted as independent findings.

### Apparatus

A custom-built NZXT desktop computer (NZXT, Los Angeles, CA, USA) equipped with MATLAB software (Mathworks, Natick, MA, USA) and Psychophysics Toolbox extensions (Brainard, 1997) was used to present the stimuli on a Sun Microsystems 4472 (Oracle Corporation, Santa Clara, CA, USA) CRT monitor (85 Hz refresh rate). The participants viewed the monitor from 70 cm in a dark, soundproof room. Eye-tracking was conducted using the EyeLink 1000 Plus system (SR Research Ltd., Ottawa, Ontario, Canada) and head position was maintained using a manufacturer-provided chin rest (SR Research Ltd.). Electric shock was administered on the third and fourth fingers using a battery powered transcutaneous aversive stimulator (Coulbourn Instruments, Allentown, PA, USA; Model Number E13-22). Electrodes were attached to the participants fingers (BIOPAC Systems, Goleta, CA, USA; EL507A) with the application of an electrode gel (BIOPAC Systems, GEL101A).

### Stimuli

For both oculomotor search paradigms, every trial comprised a gaze-contingent fixation display, a visual search array, and an inter-trial interval (ITI; see Figures 1 and 2). The fixation display comprised a fixation cross (0.7° x 0.7° visual angle) at the center of the screen. The fixation display remained on screen until eye position was registered within 1.3° of the center of the fixation cross for a continuous period of 500 ms (Kim, Senior, et al., 2024; Kim & Anderson, 2022). The visual search array was then presented for 2000 ms or until a fixation on the target was registered. If a target was not fixated within the timeout limit, the words “Eye Error” would appear in the center of the screen for 500 ms. Lastly, the ITI comprised a blank screen for 1000 ms.

**Figure 1.**
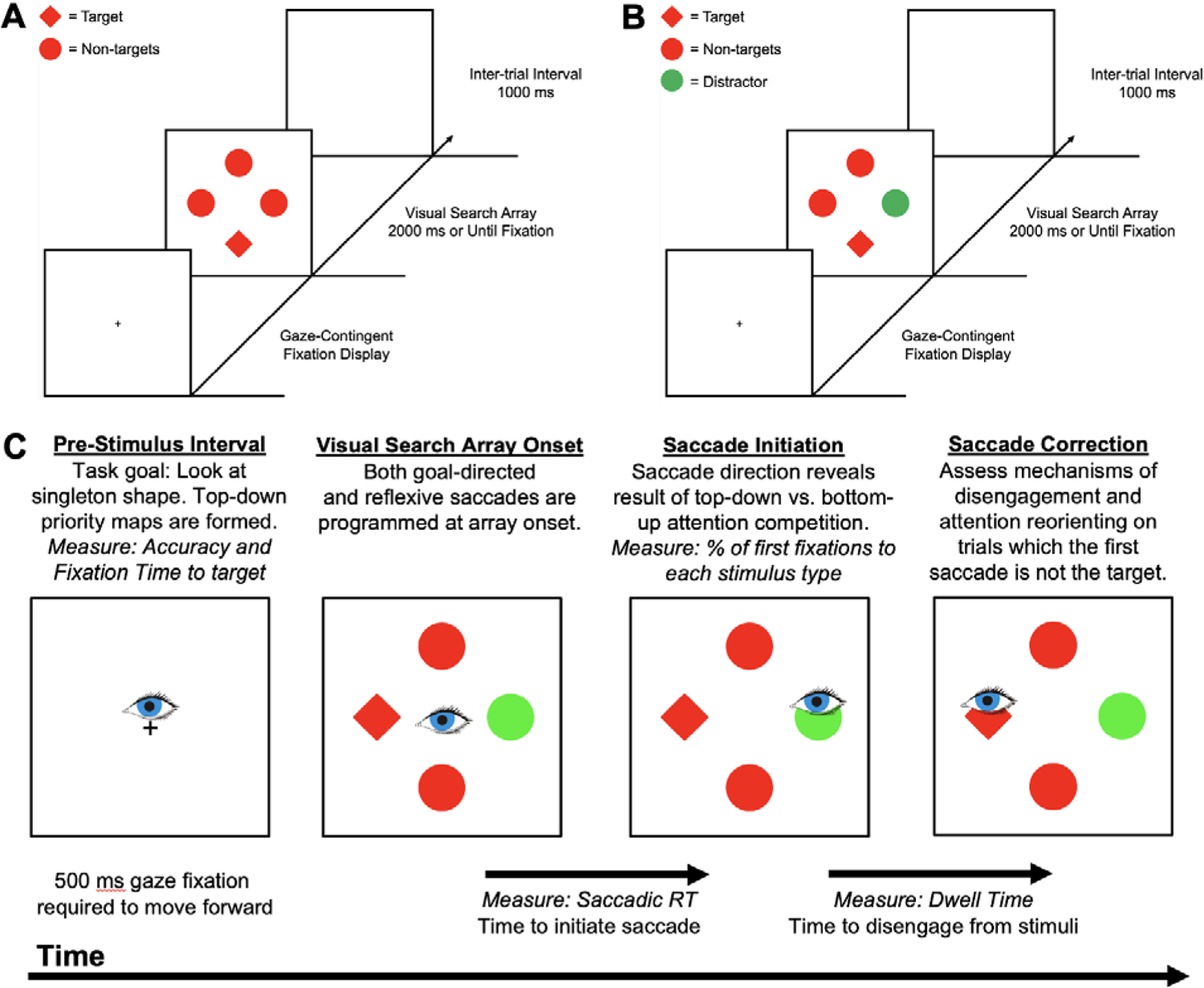
Sequence of events in the singleton-search paradigm. The singleton-search task required participants to fixate on the unique shape (circle among diamonds or diamond among circles equally often) in trials a salient distractor could be (A) absent or (B) present. (C) We illustrate the timing of individual mechanisms of attentional control including the measures acquired to assess them in a sample distractor-present trial (the eye represents gaze fixation for the purpose of the figure and is not a stimulus visible to participants).

**Figure 2.**
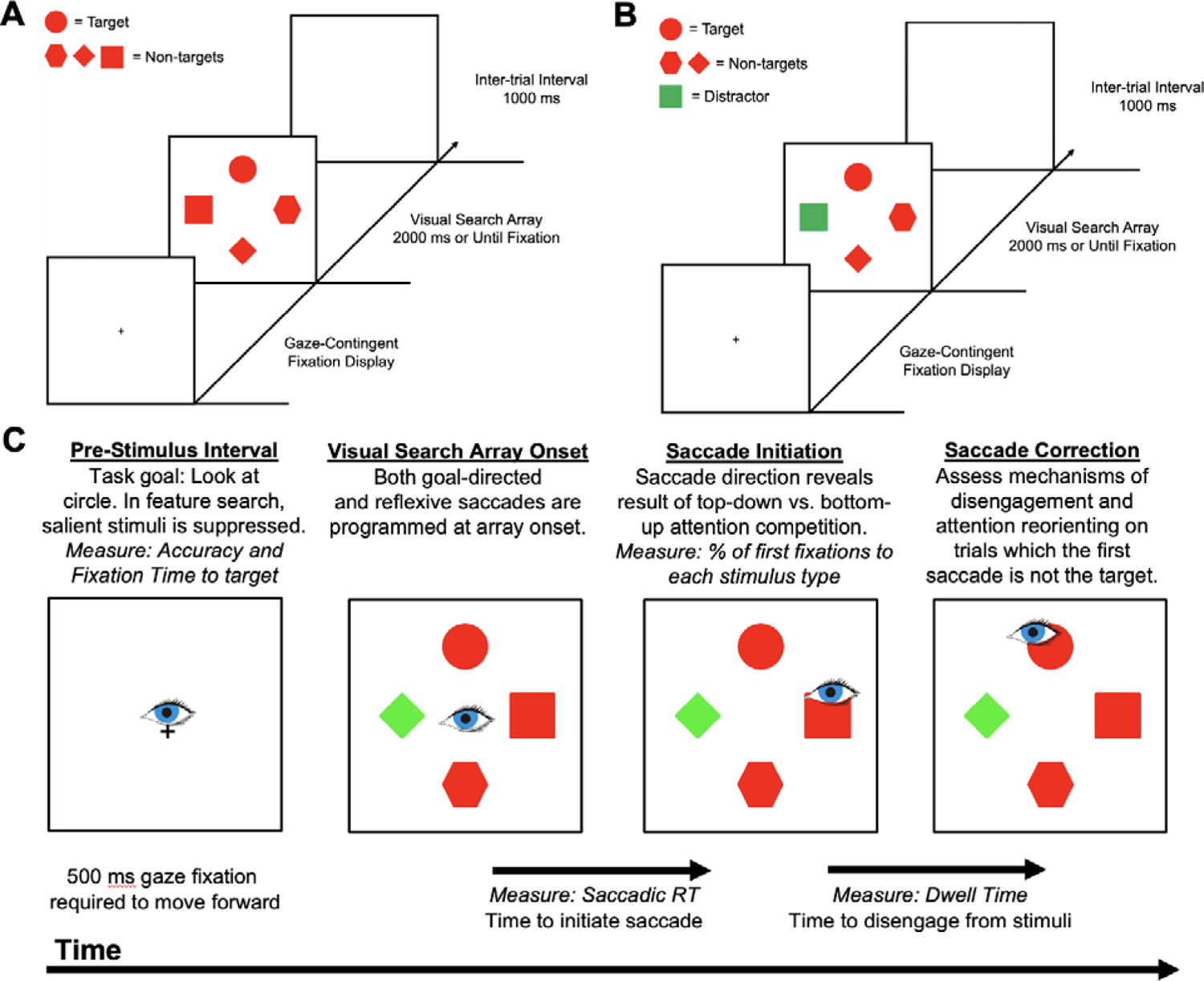
Sequence of events in feature-search paradigm. The feature-search task required participants to fixate on a specific shape (circle or diamond, counterbalanced across subjects) in trials a salient distractor could be (A) absent or (B) present. While maintaining stimuli and other design elements between experiments, this simple change in visual search array caused participants to reactively disengage from the salient distractor when engaged in singleton-search mode and proactively suppress the salient distractor when engaged in feature-search mode. (C) We illustrate the timing of individual mechanisms of attentional control including the measures acquired to assess them in a sample distractor-present trial. The eye represents gaze fixation.

In the first visual search paradigm requiring singleton search, participants had to search for the unique shape among four stimuli (one circle among three diamonds or one diamond among three circles; see Figure 1). In the second visual search paradigm requiring feature search, the search array comprised four unique shapes: a circle, diamond, square, and a hexagon. In this visual search paradigm, participants had to find a specific target shape (circle or diamond, counterbalanced across all participants; see Figure 2). For both visual search tasks, no motor responses were required and oculomotor fixation on the target ended the trial immediately. The length and width of each of the shapes was 4.2° (visual angle), except for the circle with diameter 4.0° for all shapes to have approximately equal area. Each shape was placed at equal intervals along an imaginary circle with a radius of 9.1°. The color of the shapes was red or green. In addition, the color of the distractor on distractor-present trials was counterbalanced across all participants (red distractor among green shapes or green distractor among red shapes).

The simple change in visual search array stimuli and task instructions between the two oculomotor search tasks causes participants to adopt a different attentional template and leads to differential processing of the salient, task-irrelevant distractor (see Figures 1 and 2). During singleton search, participants must assess the entire search array to find the unique shape, leading to reliable attention capture by the salient distractor and requiring reactive disengagement to reorient to the target (Kim, Senior, et al., 2024). During feature search, participants can adopt an attentional template of a specific target feature. When participants learn that the salient color distractor can never be the target (Chang & Egeth, 2021; Gaspelin et al., 2017; Stilwell & Vecera, 2022; Won & Geng, 2018), they can proactively suppress the distractor and make fewer first fixations toward the distractor, demonstrating oculomotor suppression.

### Design and Procedure

Participants first calibrated the intensity of the electric shock to an equal subjective intensity that is “unpleasant, but not painful” as in prior studies (Kim & Anderson, 2020b, 2020a). Subjects were only connected to the shock device during this calibration process or when completing the shock block to allow for the arousal-inducing nature of the device to dissipate (Kim & Anderson, 2020b). The calibration procedure started at the lowest intensity (“feels like a tickle on your finger”) and the intensity was increased step wise until the participant informed the experimenter that the intensity had become “unpleasant, but not painful.” After calibration, participants completed 20 practice trials of the task they were going to complete, ensuring they understood the task’s instructions. Participants had to reach a passing score of 90% accuracy to move forward, and the practice task would repeat until they achieved this threshold. Participants then completed two blocks of the task: no shock block and shock block. Each block comprised 4 runs of 96 trials each, totaling 384 trials. The order of whether participants completed the shock block first or second was counterbalanced across participants.

On distractor-absent trials, the location of the target was fully counterbalanced. In distractor-present trials, the location of the target and the location of the distractor in relation to the target were fully counterbalanced. Trials were randomized across all participants. In the visual search paradigm requiring singleton search, trials were distractor absent (no salient color distractor) or distractor present (one salient color distractor) equally often. The target shape also was the circle or diamond equally often and the distractor color (red or green) was counterbalanced across participants. In the visual search paradigm requiring feature search, trials were distractor absent (one target, three non-target stimuli) or distractor present (one target, two non-target stimuli, one distractor) equally often. The target shape (circle or diamond) and the distractor color (red or green) were counterbalanced across participants.

### Data Analysis

Eye position was calibrated to each run using 5-point calibration and was manually drift corrected online by the experimenter as necessary during the gaze-contingent fixation display (Kim, Senior, et al., 2024; Kim & Anderson, 2022). During the experiment, the X and Y positions of the eyes were continuously monitored in real time regarding the four stimulus positions and fixations were recorded during the experiment (fixations to every stimulus on each trial were logged and output in an excel file at the end of each run; as in, Kim, Senior, et al., 2024; Kim & Anderson, 2020b, 2020a, 2022). The EyeLink 1000 Plus also exported an EDF file after each run that contained both sample information collected at 1000 Hz and manually encoded event time stamps: presentation of the fixation stimulus, onset of the visual search array, fixation of each stimulus during visual search, and the end of the trial. Fixation of a stimulus was registered if eye position remained within a region extending 1.0° around the stimulus for a continuous period of at least 50 ms for non-targets and distractors and at least 100 ms for the target (Kim & Anderson, 2022). Saccades were defined as a minimum eye velocity threshold of 30° per second and a minimum acceleration threshold of 9,500° per second.

Fixation times were defined as the time measured from the onset of the stimulus array until a valid fixation was registered for the target shape. Fixation times that exceeded three standard deviations of the mean for a condition for each participant were trimmed (Kim, Senior, et al., 2024; Kim & Anderson, 2022). First fixations were defined as the first stimulus that was initially registered as a fixation on each trial. First fixations to the target, non-target stimuli, and distractor stimuli, in addition to accuracy and fixation times, were derived from the eye data recorded online during the experiment (see publicly available oculomotor data files in the Open Science Framework below). Percent of saccades to the non-target stimuli were corrected to provide a per-item estimate of fixation; on distractor-absent and distractor-present trials, the percentage of saccades was divided by 3 and 2, respectively. Dwell times were defined as the duration that the eyes remained within the fixation window of the stimulus and were also recorded online. Saccadic reaction times (sRT) were computed as the time to initiate a saccade relative to the onset of the visual search array (Kim, Senior, et al., 2024; Kim & Anderson, 2022). Mean sRTs were computed offline from the EyeLink 1000 EDF output files using the edfmex MATLAB executable file and the ‘STARTSACC’ code string from the event structure.

Statistical analyses were completed using SPSS (IBM, Armonk, NY, USA). Cohen’s *d* measures of effect sizes were calculated using corrected standard deviations of the mean difference as output by SPSS. Mixed analysis of variance (MANOVA) analyses were conducted with age as a between-subjects factor and all other measures as a within-subjects measure. Post-hoc *t-*test analyses were completed using paired samples comparisons for within-subjects measures and independent sample comparisons for between-subjects measures.

### Data Availability

The data have been made publicly available on the Open Science Framework, https://osf.io/hmfzq/.

## Results

### Age Differences in Task Performance during Singleton Search

First, we investigated whether the threat of unpredictable shock had a differential effect on mean accuracy (whether a fixation was made on the target within the time limit) between young and older adults. We conducted a 2×2×2 MANOVA analysis over factors age (young vs. older), distractor presence (absent vs. present), and arousal (no shock vs. shock) and found a significant main effect of age, *F*(1,66) = 5.88, *p* = .018, η*_p_ ^2^* = .082, and distractor presence, *F*(1,66) = 10.08, *p* = .002, η*_p_^2^* = .132, but no main effect of arousal, *F*(1,66) = 1.73, *p* = .193, nor any interaction effects, *Fs*(1,66) < 1.63, *ps* > .265 (see Figure 3A). When conducting the aforementioned MANOVA analysis with fixation time as the dependent variable, we found a significant main effect of age, *F*(1,66) = 31.68, *p* < .001, η*_p_^2^* = .324, distractor presence, *F*(1,66) = 345.82, *p* < .001, η*_p_^2^* = .840, but again no main effect of arousal, *F*(1,66) = .40, *p* = .528. In addition, we found a significant age x distractor presence interaction effect, *F*(1,66) = 28.86, *p* < .001, η*_p_^2^* = .304, but no other interaction effects, *Fs*(1,66) < 1.05, *ps* > .309 (see Figure 3B). Post-hoc *t-test* comparisons revealed that older adults were slower in fixating the target compared with young adults and that fixation times on distractor-present trials were longer than on distractor-absent trials for both age groups (see Table 1). The significant age x distractor presence interaction effect shows that the distractor-presence cost was greater in older adults, that is older adults were more negatively affected by the distractor with slower fixation times compared with young adults. This distractor-presence cost was evident even when normalizing fixation time speeds across age groups (see Kim, Senior, et al., 2024).

**Figure 3.**
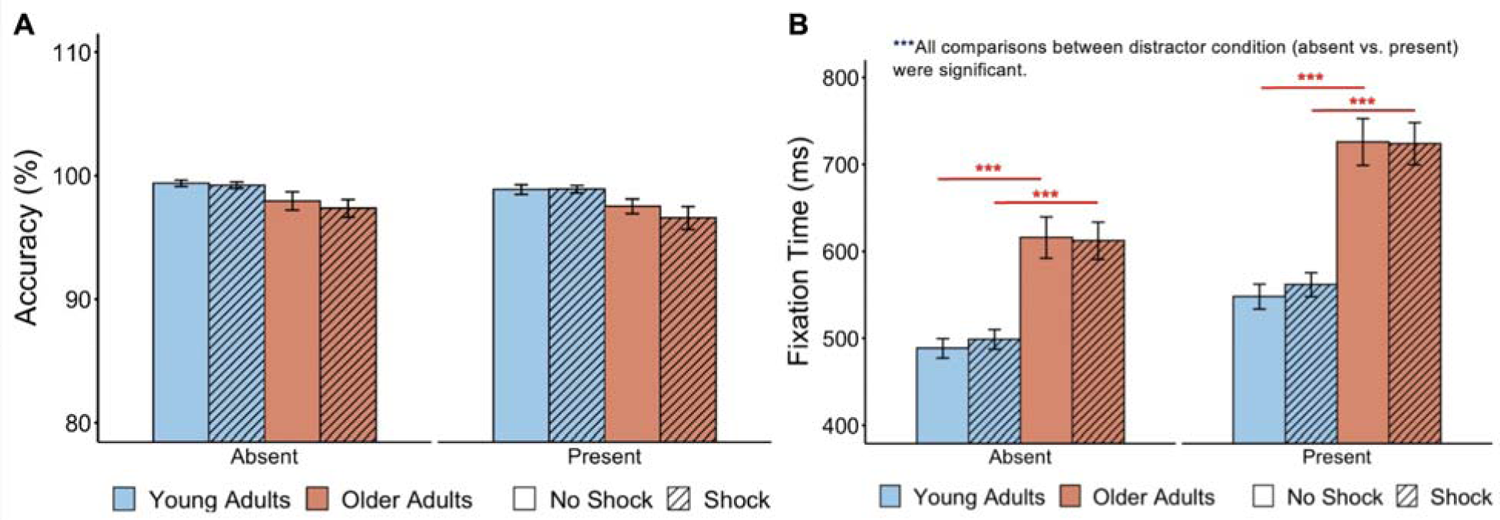
Threat of unpredictable shock did not modulate accuracy or fixation times while in singleton-search mode. Bar graphs depict age differences in (A) accuracy and (B) fixation time on distractor-absent and distractor-present trials while in singleton-search mode. The presence of the distractor decreased accuracy and slowed fixation times. Older adults also had slower fixation times compared with young adults (see red asterisks). Error bars reflect the standard error of the mean. \**\*\*p < .*001.

**Table 1.**
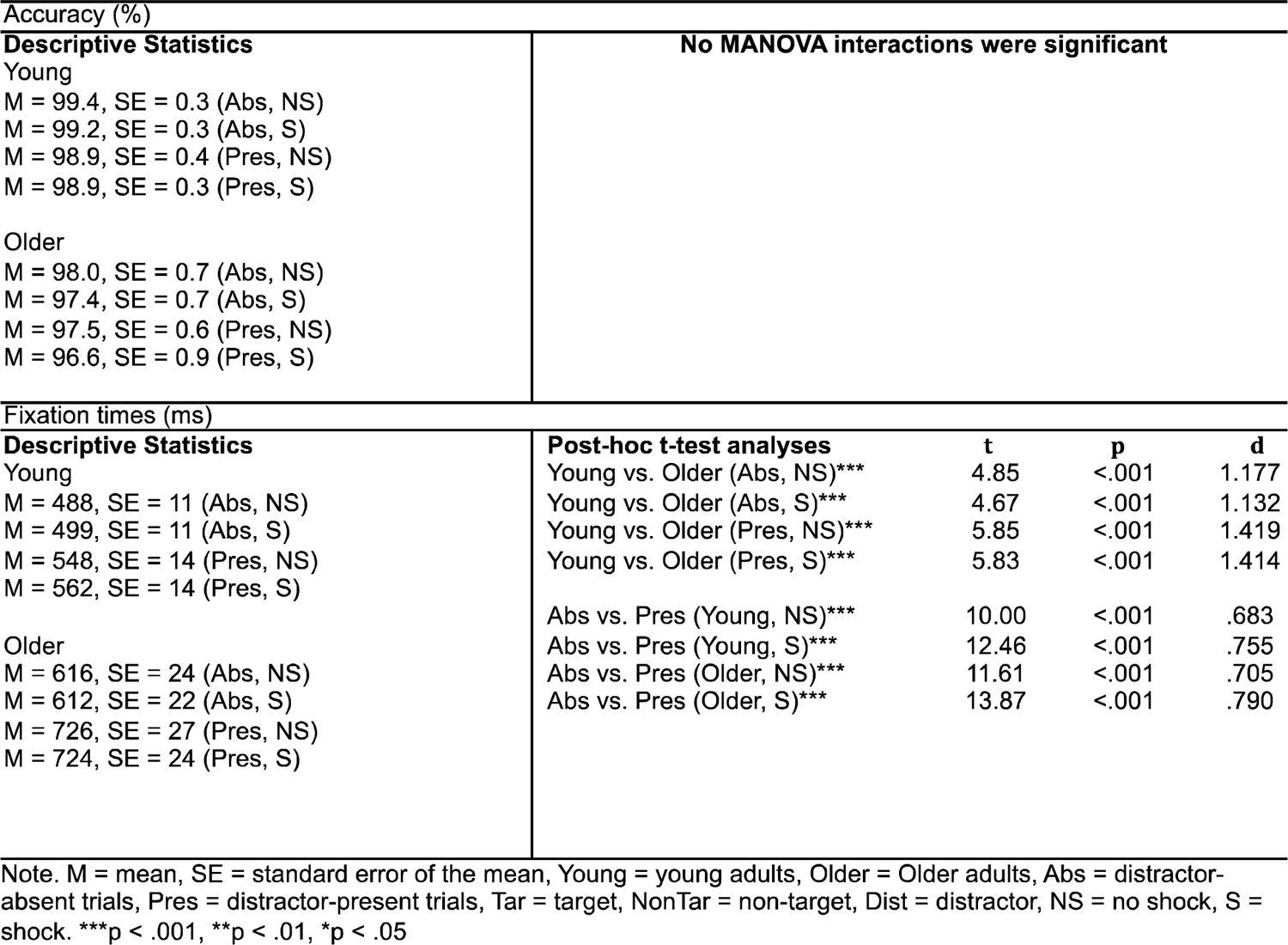
Descriptive statistics and post-hoc t-test analyses during singleton search for dependent measures accuracy and fixation time.

### Age Differences in First-Saccade Destinations during Singleton Search

First-saccade destinations reveal whether a goal-directed or reflexive saccade was initiated following competition between top-down and bottom-up networks of attention (Corbetta & Shulman, 2002). Both goal-directed and reflexive saccades are programmed simultaneously at the onset of the visual search array (Case & Ferrera, 2007; Nummela & Krauzlis, 2011; Pierrot-Deseilligny et al., 1995; Schall, 1995; Theeuwes et al., 1998). A first saccade toward the target would reveal a goal-directed saccade was initiated while a first saccade toward other stimuli would show a reflexive saccade was initiated following interactions between multiple networks of attention and visual attention priority maps (Itti & Koch, 2000, 2001). Greater first saccades toward the salient distractor compared to the average of non-target stimuli would show oculomotor capture by the salient distractor.

We first conducted a 2×2×2 MANOVA analysis over factors age (young vs. older), first-saccade destination (target vs. non-targets), and arousal (no shock vs. shock) on distractor-absent trials. Distractor-absent trials allow for a more accurate assessment of mechanisms of goal-directed attentional control without interactions of bottom-up attentional control from the visual cortex induced by a physically salient stimulus (Itti & Koch, 2001). We found a significant main effect of age, *F*(1,66) = 11.34, *p* < .001, η*_p_^2^* = .147, first-saccade destination, *F*(1,66) = 1760.18, *p* < .001, η*_p_ ^2^* = .964, but no main effect of arousal, *F*(1,66) = .20, *p* = .660 (see Figure 4A). We also found an age x first-saccade destination interaction effect, *F*(1,66) = 11.34, *p* < .001, η*_p_^2^* = .147, but did not find any other interaction effects, *Fs*(1,66) < .20, *p*s > .660. Post-hoc analyses revealed that both age groups made significantly more saccades toward the target compared with non-target stimuli, but older adults make significantly fewer first saccades toward the target and more saccades toward a non-target shape compared with young adults, demonstrating age-related impairments in goal-directed attentional control (see Table 2). Interestingly, we observed that arousal did not modulate goal-directed attentional control in both age groups.

**Figure 4.**
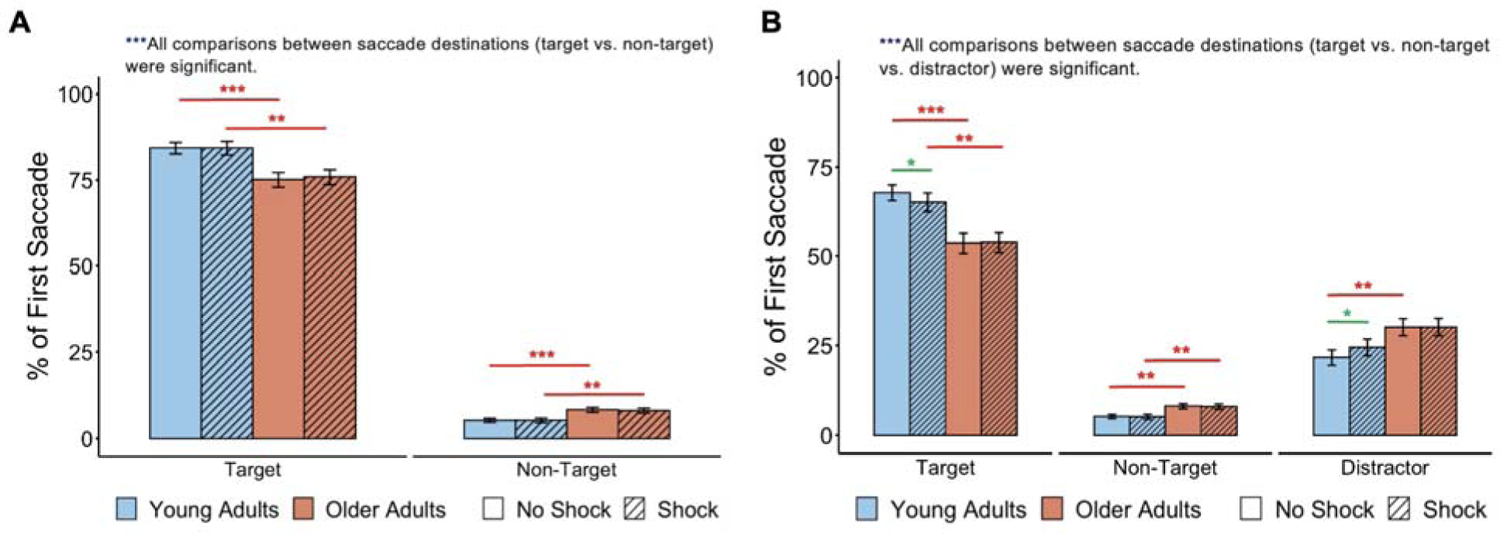
Threat of shock increased distractibility by salient distractors in young adults but not in older adults while in singleton-search mode. Bar graphs show first-saccade destinations on (A) distractor-absent and (B) distractor-present trials. Both age groups made more first saccades toward the target compared to non-target shapes showing goal-directed attentional control and more first saccades toward the distractor compared to non-target shapes showing attention capture by the distractor (see also Table 2). However, older adults had made fewer first saccades toward the target than did young adults, indicating age-related impairments in goal-directed attentional control (see red asterisks). The threat of unpredictable shock increased distractibility by salient distractors in young adults (see green asterisks), but not in older adults. Error bars reflect the standard error of the mean. *\*\*\*p < .*001*, **p < .*01*, *p < .*05.

**Table 2.** Descriptive statistics and post-hoc t-test analyses during singleton search for dependent measures first-saccade destination.

| First-saccade destination on distractor-absent trials (%) |  |  |  |  |
| --- | --- | --- | --- | --- |
| Descriptive Statistics | Post-hoc t-test analyses | t | p | d |
| Young | Young vs. Older (Tar, NS)*** | 3.50 | <.001 | .848 |
| M = 84.3, SE = 1.6 (Tar, NS) | Young vs. Older (Tar, S)** | 2.93 | .005 | .711 |
| M = 84.3, SE = 2.0 (Tar, S) | Young vs. Older (NonTar, NS)*** | 3.50 | <.001 | .848 |
| M = 5.2, SE = 0.5 (NonTar, NS) | Young vs. Older (NonTar, S)** | 2.93 | .005 | .711 |
| M = 5.2, SE = 0.7 (NonTar, S) |  |  |  |  |
| Older | Tar vs. NonTar (Young, NS)*** | 36.07 | <.001 | 12.372 |
| M = 75.1, SE = 2.1 (Tar, NS) | Tar vs. NonTar (Young, S)*** | 30.30 | <.001 | 10.392 |
| M = 75.8, SE = 2.1 (Tar, S) | Tar vs. NonTar (Older, NS)*** | 24.41 | <.001 | 8.373 |
| M = 8.3, SE = 0.7 (NonTar, NS) | Tar vs. NonTar (Older, S)*** | 24.15 | <.001 | 8.285 |
| M = 8.1, SE = 0.7 (NonTar, S) |  |  |  |  |
| First-saccade destination on distractor-present trials (%) |  |  |  |  |
| Descriptive Statistics | Post-hoc t-test analyses | t | p | d |
| Young | Young vs. Older (Tar, NS)*** | 3.96 | <.001 | .961 |
| M = 67.9, SE = 2.2 (Tar, NS) | Young vs. Older (Tar, S)** | 2.93 | .005 | .711 |
| M = 65.3, SE = 2.7 (Tar, S) | Young vs. Older (NonTar, NS)** | 3.26 | .002 | .791 |
| M = 5.2, SE = 0.5 (NonTar, NS) | Young vs. Older (NonTar, S)** | 2.78 | .007 | .674 |
| M = 5.1, SE = 0.7 (NonTar, S) | Young vs. Older (Dist, NS)** | 2.69 | .009 | .653 |
| M = 21.6, SE = 2.1 (Dist, NS) | Young vs. Older (Dist, S) | 1.69 | .097 |  |
| M = 24.5, SE = 2.4 (Dist, S) |  |  |  |  |
| Older | Tar vs. NonTar (Young, NS)*** | 25.64 | <.001 | 7.208 |
| M = 53.6, SE = 2.9 (Tar, NS) | NonTar vs. Dist (Young, NS)*** | 7.31 | <.001 | 1.907 |
| M = 53.9, SE = 2.8 (Tar, S) | Tar vs. Dist (Young, NS)*** | 11.12 | <.001 | 3.690 |
| M = 8.1, SE = 0.7 (NonTar, NS) | Tar vs. NonTar (Young, S)*** | 19.78 | <.001 | 5.808 |
| M = 8.0, SE = 0.8 (NonTar, S) | NonTar vs. Dist (Young, S)*** | 7.75 | <.001 | 1.934 |
| M = 30.1, SE = 2.4 (Dist, NS) | Tar vs. Dist (Young, S)*** | 8.44 | <.001 | 2.790 |
| M = 30.2, SE = 2.4 (Dist, S) | Tar vs. NonTar (Older, NS)*** | 13.72 | <.001 | 4.172 |
|  | NonTar vs. Dist (Older, NS)*** | 9.19 | <.001 | 2.119 |
|  | Tar vs. Dist (Older, NS)*** | 4.65 | <.001 | 1.542 |
|  | Tar vs. NonTar (Older, S)*** | 13.92 | <.001 | 4.154 |
|  | NonTar vs. Dist (Older, S)*** | 4.68 | <.001 | 1.540 |
|  | Tar vs. Dist (Older, S)*** | 8.64 | <.001 | 2.121 |
|  | NS vs. S (Young, Tar)* | 2.75 | .010 | .168 |
|  | NS vs. S (Young, NonTar) | .18 | .861 |  |
|  | NS vs. S (Young, Dist)* | 2.58 | .015 | .211 |
|  | NS vs. S (Older, Tar) | .15 | .885 |  |
|  | NS vs. S (Older, NonTar) | .32 | .749 |  |
|  | NS vs. S (Older, Dist) | .02 | .981 |  |

Note. M = mean, SE = standard error of the mean, Young = young adults, Older = Older adults, Abs = distractor-absent trials, Pres = distractor-present trials, Tar = target, NonTar = non-target, Dist = distractor, NS = no shock, S = shock. \*\*\* $p < .001$ , \*\* $p < .01$ , \* $p < .05$

In distractor-present trials, we can determine that the highest saliency item in the visual search array captured attention if the first-saccade destination is a reflexive saccade toward the distractor. If more reflexive first saccades go to the distractor compared to a non-target showing an oculomotor capture effect, we can quantify the magnitude of attentional priority allocation toward the salient color stimuli. We conducted a 2×3×2 MANOVA analysis over factors age (young vs. older), first-saccade destination (target vs. non-targets vs. distractor), and arousal (no shock vs. shock) on distractor-present trials. We observed a significant main effect of age, *F*(1,66) = 9.81, *p* = .003, η*_p_^2^* = .129, first-saccade destination, *F*(2,132) = 245.37, *p* < .001, η*_p_^2^* = .788, but no main effect of arousal, *F*(1,66) = .13, *p* = .725 (see Figure 4A). However, we observed an age x first-saccade destination interaction effect, *F*(2,132) = 9.27, *p* < .001, η*_p_^2^* = .123 and a trending three-way interaction between factors age, arousal, and first-saccade destination, *F*(2,132) = 2.62, *p* = .076. Given that our primary research question was to investigate whether increased arousal would differentially modulate attention capture by the salient distractor in young and older adults, we further explored this three-way interaction by conducting a 2×2 MANOVA with factors age (young vs. older) and arousal (no shock vs. shock) on first-saccade destinations to the salient distractor. We again observed a marginal interaction effect, *F*(1,66) = 3.00, *p* = .088. Post-hoc *t-*tests identified that young adults made significantly more first-saccades toward the distractor, *t*(33) = 2.58, *p* = .015, *d* = .211, as previously observed in this visual search paradigm under threat of unpredictable shock (Kim & Anderson, 2020b). However, post-hoc *t-*tests revealed no evidence that older adults were more distractible by salient stimuli under increased arousal, *t*(33) = .02, *p =* .981 (see also Table 2). Given the marginal MANOVA interaction and the null findings in older adults, we conducted a Bayesian paired samples t-test using JASP (ver 0.18.3) to investigate the evidence in support of the null hypothesis in older adults, H_0_ = arousal does not modulate attention capture by the salient distractor (Wagenmakers, Love, et al., 2018; Wagenmakers, Marsman, et al., 2018). The Bayes factor (BF_10_) measures the strength of evidence in support of the alternative hypothesis compared to the null hypothesis H_0_ and a BF_10_ value of 1 to .33 reflects anecdotal evidence, .33 to .1 moderate evidence, and < .1 as strong evidence in favor of the null hypothesis. We found moderate evidence that arousal does not modulate attention capture by the salient distractor in older adults, BF_10_ = .184 (error% = 0.041).

Importantly, we found that under the threat of unpredictable shock, young adults made fewer first saccades to the target and more first saccades to the distractor, demonstrating increased distractibility by salient stimuli during states of increased arousal (see Table 2). We did not observe this effect of arousal-biased competition in older adults. Furthermore, the decreased frequency of first saccades to the target did not lead to a general increase in reflexive saccades in young adults. Instead, the specific increase in attentional priority of the salient stimulus in the search array during heightened states of arousal was only observed through an increase in first saccades toward the salient distractor. The decrease in first saccades toward the target under arousal appeared to result from increased first saccades to distracters reducing first saccades to targets, as we did not observe this arousal-induced pattern in distractor-absent trials.

### Age Differences in Dwell Time and Saccadic Reactions Times during Singleton Search

We investigated how long participants fixated on an incorrect initial saccade (when the first-saccade destination is not the target), or dwell time, as a measure of reactive disengagement and reorienting following an incorrect fixation. Longer dwell times would show a prolonged processing requirement of a stimulus until the next saccade is initiated and shows deficiencies in rejecting a stimulus as a non-target stimulus. A 2×2×2 MANOVA analysis over factors age (young vs. older), stimuli (non-targets vs. distractors), and arousal (no shock vs. shock) revealed a significant main effect of age, *F*(1,66) = 13.18, *p* < .001, η*_p_^2^* = .166, stimuli, *F*(1,66) = 51.77, *p* < .001, η*_p_^2^* = .440, but no main effect of arousal, *F*(1,66) = .89, *p* = .350 (see Figure 5A). We also observed a significant age x stimuli interaction effect, *F*(1,66) = 4.67, *p* = .034, η*_p_^2^* = .066 but no other interaction effects, *Fs*(1,66) < .51, *ps* > .477. Post-hoc analyses revealed that both young and older adults had longer fixation durations on the distractor compared to a non-target shape and that older adults had longer fixation durations compared with young adults (see Table 3). Furthermore, the significant interaction effect shows that older adults had particularly longer dwell times compared with young adults when the stimulus was the distractor, demonstrating an age-related impairment in disengaging from salient stimuli compared with non-salient stimuli.

**Figure 5.**
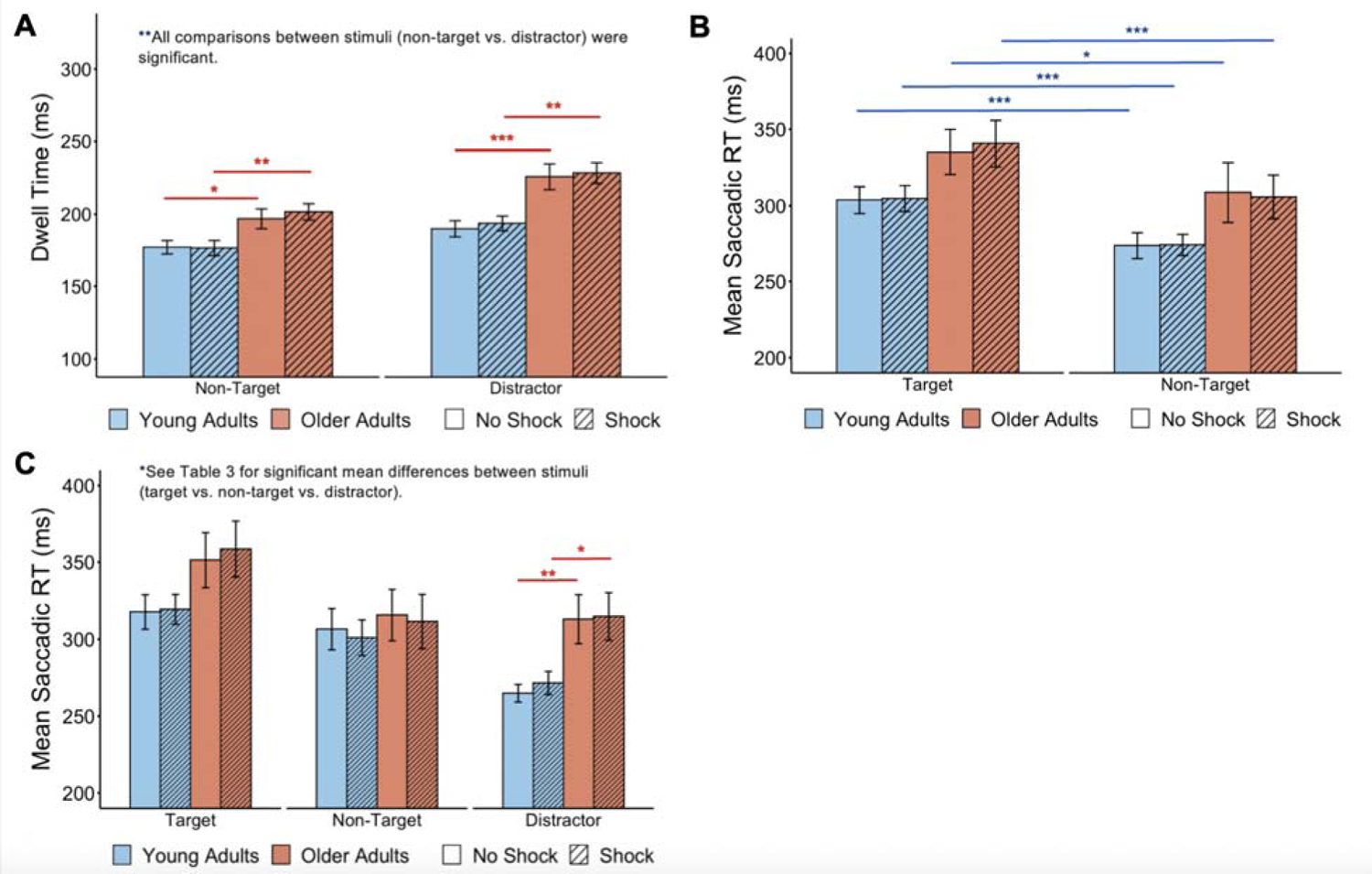
Threat of shock did not modulate dwell time or saccade latencies while in singleton-search mode. Bar graphs depict (A) dwell time and mean saccadic reaction times in (B) distractor-absent and (C) distractor-present trials. Both age groups had longer fixation durations on the distractor compared to non-target shapes and had slower mean saccade latencies when making goal-directed saccades toward the target (blue asterisks; see also Table 3). However, older adults showed significantly longer dwell times and longer mean saccade latencies when the direction of the saccade was the distractor (see red asterisks). Error bars reflect standard error of the mean. \**p* < .05, \*\**p* < .01, \*\*\**p* < .001.

**Table 3.** Descriptive statistics and post-hoc t-test analyses during singleton search for dependent measures dwell time and mean saccadic reaction time.

| Dwell time (ms) |  |  |  |  |
| --- | --- | --- | --- | --- |
| Descriptive Statistics | Post-hoc t-test analyses | t | p | d |
| Young | Young vs. Older (NonTar, NS)* | 2.39 | .020 | .581 |
| M = 177.0, SE = 4.7 (NonTar, NS) | Young vs. Older (NonTar, S)** | 2.720 | .008 | .660 |
| M = 176.5, SE = 5.3 (NonTar, S) | Young vs. Older (Dist, NS)*** | 3.43 | <.001 | .832 |
| M = 190.0, SE = 5.6 (Dist, NS) | Young vs. Older (Dist, S)** | 3.959 | .005 | .960 |
| M = 193.7, SE = 5.1 (Dist, NS) |  |  |  |  |
| Older | NonTar vs. Dist (Young, NS)** | 3.17 | .003 | .421 |
| M = 196.9, SE = 6.8 (NonTar, NS) | NonTar vs. Dist (Young, S)*** | 3.90 | <.001 | .566 |
| M = 201.6, SE = 7.6 (NonTar, S) | NonTar vs. Dist (Older, NS)*** | 4.76 | <.001 | .604 |
| M = 225.9, SE = 8.9 (Dist, NS) | NonTar vs. Dist (Older, S)*** | 4.35 | <.001 | .624 |
| M = 228.5, SE = 7.2 (Dist, S) |  |  |  |  |
| Mean saccadic reaction times on distractor-absent trials (ms) |  |  |  |  |
| Descriptive Statistics | No MANOVA interactions were significant |  |  |  |
| Young |  |  |  |  |
| M = 303.7, SE = 8.9 (Tar, NS) |  |  |  |  |
| M = 304.8, SE = 8.6 (Tar, S) |  |  |  |  |
| M = 273.7, SE = 8.6 (NonTar, NS) |  |  |  |  |
| M = 274.1, SE = 7.1 (NonTar, S) |  |  |  |  |
| Older |  |  |  |  |
| M = 335.3, SE = 14.6 (Tar, NS) |  |  |  |  |
| M = 340.8, SE = 15.2 (Tar, S) |  |  |  |  |
| M = 308.8, SE = 19.7 (NonTar, NS) |  |  |  |  |
| M = 305.9, SE = 14.3 (NonTar, S) |  |  |  |  |
| Mean saccadic reaction times on distractor-present trials (ms) |  |  |  |  |
| Descriptive Statistics | Post-hoc t-test analyses | t | p | d |
| M = 317.7, SE = 11.2 (Tar, NS) | Young vs. Older (Tar, NS) | 1.60 | .115 |  |
| M = 319.4, SE = 9.7 (Tar, S) | Young vs. Older (Tar, S) | 1.88 | .064 |  |
| M = 306.5, SE = 13.4 (NonTar, NS) | Young vs. Older (NonTar, NS) | .43 | .669 |  |
| M = 301.0, SE = 11.6 (NonTar, S) | Young vs. Older (NonTar, S) | .50 | .620 |  |
| M = 264.9, SE = 5.7 (Dist, NS) | Young vs. Older (Dist, NS)** | 2.84 | .006 | .689 |
| M = 271.5, SE = 7.6 (Dist, S) | Young vs. Older (Dist, S)* | 2.51 | .015 | .608 |
| Older | Tar vs. NonTar (Young, NS) | 1.62 | .115 |  |
| M = 351.3, SE = 17.8 (Tar, NS) | NonTar vs. Dist (Young, NS)*** | 3.79 | <.001 | .582 |
| M = 358.5, SE = 18.3 (Tar, S) | Tar vs. Dist (Young, NS)*** | 6.47 | <.001 | .835 |
| M = 315.7, SE = 16.7 (NonTar, NS) | Tar vs. NonTar (Young, S)** | 3.00 | .005 | .284 |
| M = 311.5, SE = 17.7 (NonTar, S) | NonTar vs. Dist (Young, S)*** | 7.91 | <.001 | .894 |
| M = 313.0, SE = 15.9 (Dist, NS) | Tar vs. Dist (Young, S)*** | 3.47 | <.001 | .476 |
| M = 314.8, SE = 15.5 (Dist, S) | Tar vs. NonTar (Older, NS)*** | 4.14 | <.001 | .352 |
|  | NonTar vs. Dist (Older, NS) | .28 | .780 |  |
|  | Tar vs. Dist (Older, NS)*** | 6.44 | <.001 | .371 |
|  | Tar vs. NonTar (Older, S)*** | 4.68 | <.001 | .447 |
|  | NonTar vs. Dist (Older, S) | .34 | .735 |  |
|  | Tar vs. Dist (Older, S)*** | 7.21 | <.001 | .393 |

Note. M = mean, SE = standard error of the mean, Young = young adults, Older = Older adults, Abs = distractor-absent trials, Pres = distractor-present trials, Tar = target, NonTar = non-target, Dist = distractor, NS = no shock, S = shock. \*\*\*p < .001. \*\*p < .01, \*p < .05

Next, we investigated whether the threat of unpredictable shock produced age differences in mean saccadic reaction times (mean sRT; also known as saccadic latencies) or the time to initiate the first saccade from the onset of the visual search array. Mean sRTs measure the time required to resolve competition between top-down and bottom-up networks of attention and determine how long it takes to initiate a reflexive saccade or to initiate a goal-directed saccade after rejecting the reflexive saccade. First, we conducted a 2×2×2 MANOVA analysis over factors age (young vs. older), stimuli (target vs. non-target), and arousal (no shock vs. shock) on distractor-absent trials. We found a significant main effect of age, *F*(1,66) = 4.04, *p* = .048, η*_p_^2^* = .058, stimuli, *F*(1,66) = 49.69, *p* < .001, η*_p_ ^2^* = .430, but no main effect of arousal, *F*(1,66) = .05, *p* = .816, nor any interaction effects, *Fs*(1,66) < .39, *ps* > .533 (see Figure 5B).

Mean sRTs are longer when the direction of the first saccade is toward the target because of processing of goal-directed mechanisms of attention compared with an automatic reflexive saccade driven by the visual cortex. Furthermore, we identified that older adults exhibit longer mean sRTs compared with young adults showing a longer time requirement to process visual stimuli (see Table 3). Finally, we conducted a 2×3×2 MANOVA analysis over factors age (young vs. older), stimuli (target vs. non-target vs. distractor), and arousal (no shock vs. shock) on distractor-present trials. We found a marginal main effect of age, *F*(1,66) = 2.84, *p* = .097, a significant main effect of stimuli, *F*(2,132) = 48.15, *p* < .001, η*_p_ ^2^* = .422, but no main effect of arousal, *F*(1,66) = .09, *p* = .767 (see Figure 5C). In addition, we found a significant age x stimuli interaction effect, *F*(2,132) = 7.84, *p* < .001, η*_p_^2^* = .106, but no other interaction effects, *Fs*(2,132) < 1.19, *ps* > .306. As in distractor-absent trials, mean sRTs were longer when making goal-directed saccades (toward target) compared to when making reflexive saccades and older adults had slower mean sRTs compared with young adults (see Table 3). However, older adults had significantly longer mean sRTs compared with young adults when the direction of the saccade was toward the distractor, as showed by the interaction effect.

In this next section, we present results from the second oculomotor search task in which participants searched for a specific shape while engaged in feature-search mode (see Figure 2). We were particularly interested in whether young adults would exhibit the persistence of arousal-biased competition, with increased first-saccade destinations towards the salient distractor, even when the distractor is proactively suppressed and whether older would exhibit changes in mechanisms of arousal-biased competition during feature search. Furthermore, we explored whether the effects of arousal on attention and oculomotor processing were equal for both reactive (singleton search) and proactive (feature search) mechanisms of attentional control. Divergent findings across tasks would suggest differences in the neural mechanisms in which arousal and the locus coeruleus-noradrenaline system modulate reactive vs. proactive mechanisms and potential differences in the neural networks underlying these distinct processes (Geng, 2014).

### Age Differences in Task Performance during Feature Search

We conducted a 2×2×2 MANOVA analysis over factors age (young vs. older), distractor presence (absent vs. present), and arousal (no shock vs. shock) on mean accuracy. We found a significant main effect of age, *F*(1,70) = 5.13, *p* = .027, η *^2^* = .068, and distractor presence, *F*(1,70) = 5.59, *p* = .021, η*_p_^2^* = .074, but no main effect of arousal, *F*(1,70) = 1.35, *p* = .250, nor any interaction effects, *Fs*(1,66) < 1.84, *ps* > .180 (see Figure 6A and Table 4). When conducting the aforementioned MANOVA analyses on fixation time, we found no main effects of age, *F*(1,70) = 3.13, *p* = .081, distractor presence, *F*(1,70) = .03, *p* = .866, a marginal effect of arousal, *F*(1,70) = 3.64, *p* = .060, nor any interaction effects, *Fs*(1,70) < 1.31, *ps* > .257 (see Figure 6B and Table 4). In this marginal effect of arousal, the threat of unpredictable shock led to a trend of slowing in overall performance across age groups.

**Figure 6.**
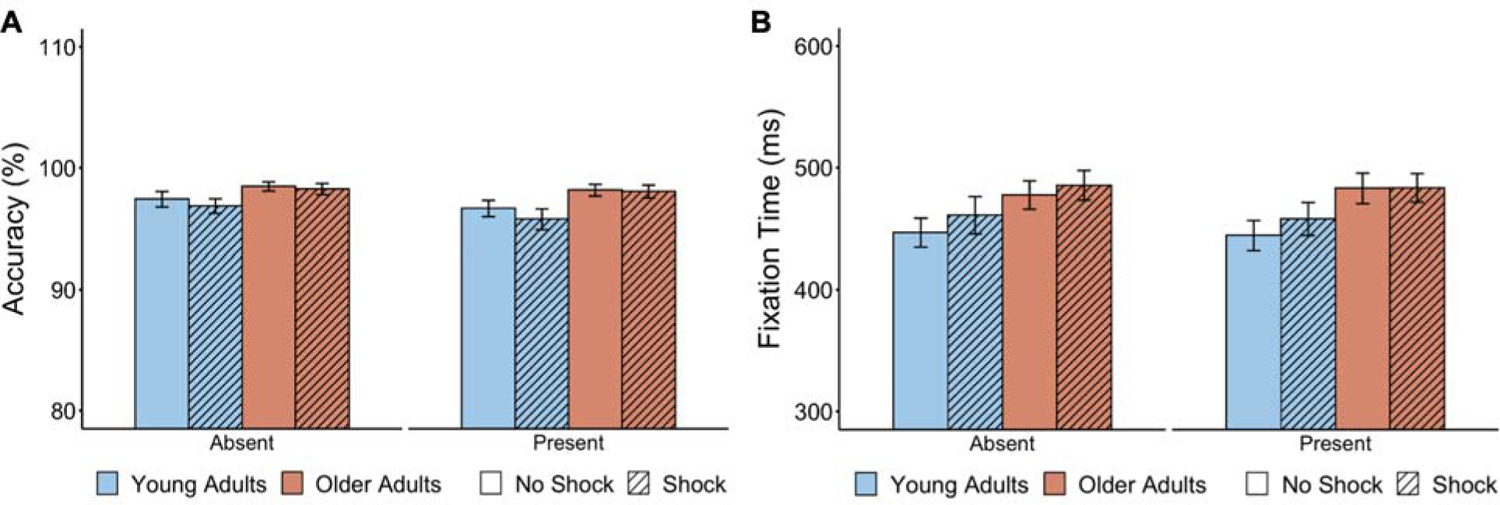
Threat of shock did not modulate accuracy and marginally slowed fixation times while in feature-search mode. Bar graphs depict age differences in (A) accuracy and (B) fixation time on distractor-absent and distractor-present trials while in feature-search mode. Error bars reflect the standard error of the mean.

**Table 4.** Descriptive statistics and post-hoc t-test analyses during feature search for dependent measures.

| Accuracy (%) |  |
| --- | --- |
| Descriptive Statistics | No MANOVA interactions were significant |
| Young<br>M = 97.4, SE = 0.6 (Abs, NS)<br>M = 96.8, SE = 0.6 (Abs, S)<br>M = 96.6, SE = 0.7 (Pres, NS)<br>M = 95.7, SE = 0.9 (Pres, S) |  |
| Older<br>M = 98.5, SE = 0.4 (Abs, NS)<br>M = 98.2, SE = 0.5 (Abs, S)<br>M = 98.2, SE = 0.5 (Pres, NS)<br>M = 98.0, SE = 0.5 (Pres, S) |  |
| Fixation times (ms) |  |
| Descriptive Statistics | No MANOVA interactions were significant |
| Young<br>M = 446, SE = 12 (Abs, NS)<br>M = 461, SE = 15 (Abs, S)<br>M = 444, SE = 12 (Pres, NS)<br>M = 458, SE = 14 (Pres, S) |  |
| Older<br>M = 477, SE = 12 (Abs, NS)<br>M = 485, SE = 12 (Abs, S)<br>M = 483, SE = 13 (Pres, NS)<br>M = 483, SE = 12 (Pres, S) |  |

Note. M = mean, SE = standard error of the mean, Young = young adults, Older = Older adults, Abs = distractor-absent trials, Pres = distractor-present trials, Tar = target, NonTar = non-target, Dist = distractor, NS = no shock, S = shock. \*\*\* $p < .001$ , \*\* $p < .01$ , \* $p < .05$

### Age Differences in First-Saccade Destination during Feature Search

We conducted a 2×2×2 MANOVA analysis over factors age (young vs. older), first-saccade destination (target vs. non-targets), and arousal (no shock vs. shock) on distractor-absent trials. We found a significant main effect of age, *F*(1,70) = 12.49, *p* < .001, η*_p_^2^* = .151, first-saccade destination, *F*(1,70) = 1181.65, *p* < .001, η *^2^* = .944, and a marginal main effect of arousal, *F*(1,70) = 3.70, *p* = .058 (see Figure 7A). We also found an age x first-saccade destination interaction effect, *F*(1,70) = 12.49, *p* < .001, η *^2^* = .151, a marginal arousal x first-saccade destination interaction effect, *F*(1,70) = 3.70, *p* = .058, but no other interaction effects, *Fs*(1,70) = .71, *p* = .404. Post-hoc analyses revealed that both age groups made significantly more saccades toward the target compared to non-target stimuli, but older adults made significantly less first saccades toward the target and more saccades toward a non-target shape compared with young adults (see Table 5).

**Figure 7.**
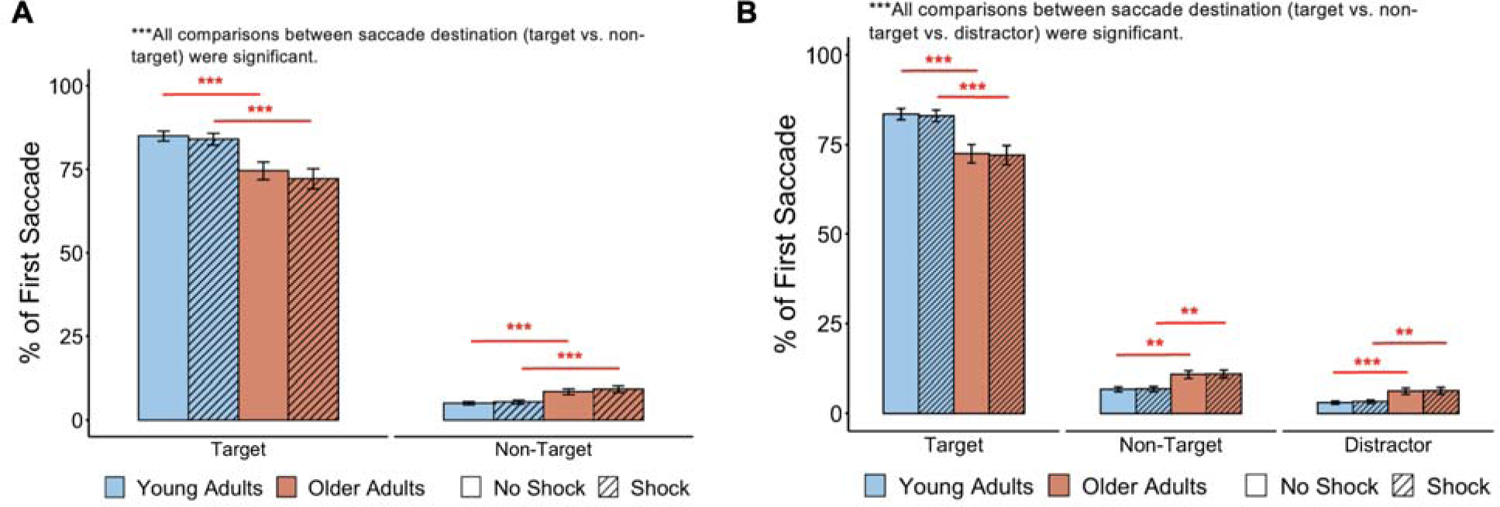
Threat of shock did not modulate first-saccade destinations while in feature-search mode. Bar graphs show first-saccade destinations on (A) distractor-absent and (B) distractor-present trials. Both age groups made more first saccades toward the target compared with non-target shapes showing goal-directed attentional control and less first saccades toward the distractor compared to non-target shapes showing an oculomotor suppression effect (see also Table 5). We found no differences in the oculomotor suppression effect between age groups. As in singleton-search mode, older adults made fewer first saccades toward the target compared with young adults showing age-related impairments in goal-directed attentional control (see red asterisks). However, unlike in singleton-search mode, the threat of unpredictable shock did not increase distractibility by salient distractors in young adults. Error bars reflect the standard error of the mean. *\*\*\*p < .001, **p < .01, *p < .05*.

**Table 5.**
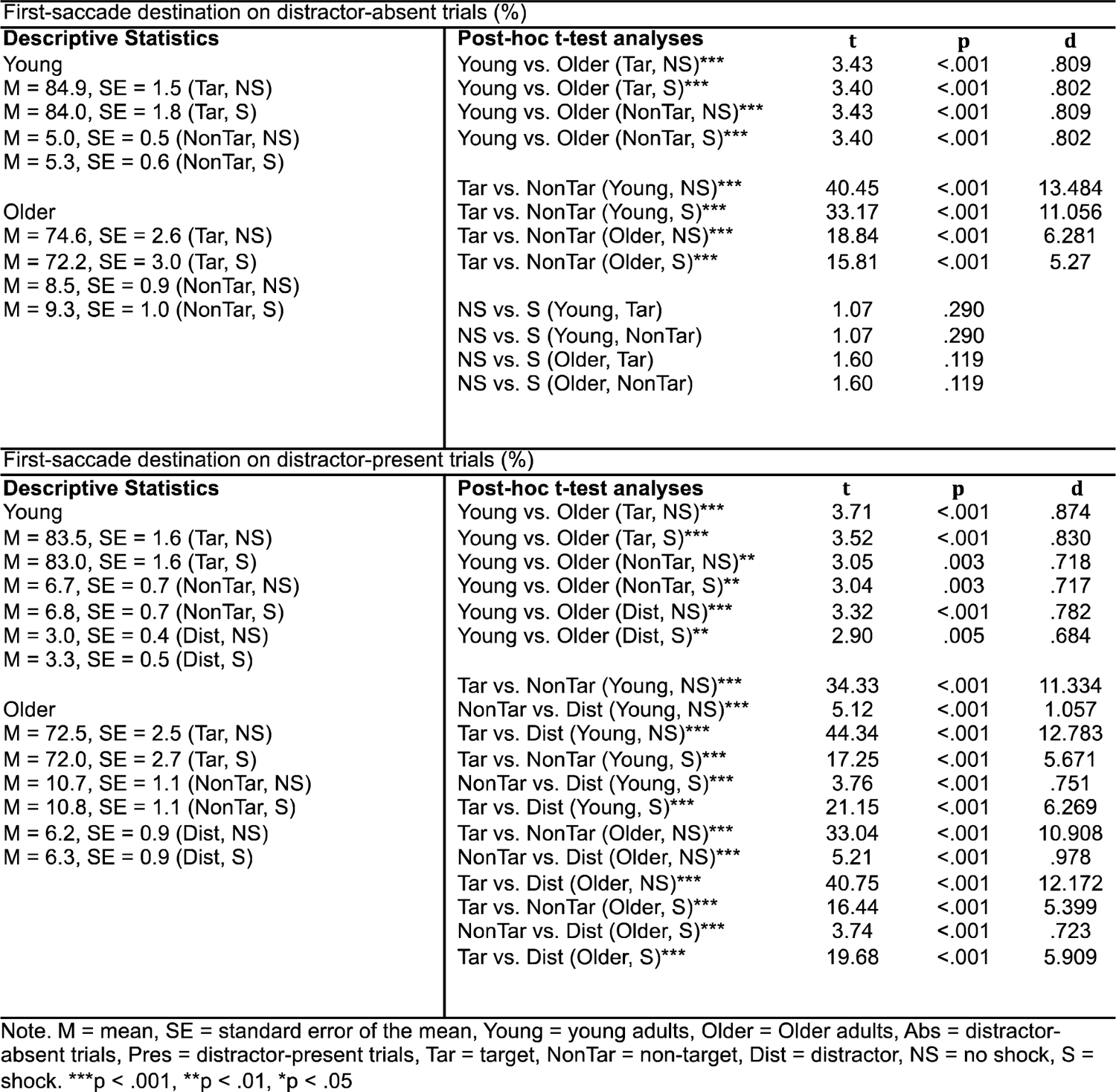
Descriptive statistics and post-hoc t-test analyses during feature search for dependent measures first-saccade destination.

For distractor-present trials, we conducted a 2×3×2 MANOVA analysis over factors age (young vs. older), first-saccade destination (target vs. non-targets vs. distractor), and arousal (no shock vs. shock). We found a significant main effect of age, *F*(1,70) = 9.89, *p* = .002, η*_p_^2^* = .124, first-saccade destination, *F*(2,140) = 1257.86, *p* < .001, η *^2^* = .947, but no main effect of arousal, *F*(1,70) = .14, *p* = .705 (see Figure 7A). We also found an age x first-saccade destination interaction effect, *F*(2,140) = 13.25, *p* < .001, η *^2^* = .159, but no other interaction effects, *Fs*(2,140) < .32, *ps* > .728. Both age groups made significantly more saccades toward the target compared with non-target stimuli and distractors and made significantly less first saccades toward the distractor compared with non-target shapes (see Table 5). Unlike in singleton-search mode, the threat of unpredictable shock did not increase attention capture by the distractor in young adults.

Lastly, we calculated the oculomotor suppression effect (% of saccades to a single non-target shape compared to a distractor) to accurately measure distractor suppression (Wöstmann et al., 2022). For young adults, the oculomotor suppression effect was 3.8% (SE = 0.3%) in no shock blocks and 3.6% (SE = 0.7%) in shock blocks. For older adults, the oculomotor suppression effect was 4.4% (SE = 1.2%) in no shock blocks and 4.5% in shock blocks (SE = 1.2%). To determine if the threat of shock differentially modulated the oculomotor suppression effect, we conducted a 2×2 ANOVA over factors age (young vs. older) and arousal (no shock vs. shock). We found no differences in the oculomotor suppression effect between young and older adults, *F*(1,70) = .38, *p =* .542, and did not observe a main effect of arousal, *F*(1,70) = .02, *p =* .885, nor an interaction effect, *F*(1,70) = .04, *p =* .835.

### Age Differences in Dwell Time and Saccadic Reactions Times during Feature Search

We conducted a 2×2×2 MANOVA analysis over factors age (young vs. older), stimuli (non-targets vs. distractors), and arousal (no shock vs. shock) over dwell time. We found a significant main effect of age, *F*(1,70) = 6.36, *p* = .014, η*_p_ ^2^* = .093, but no significant main effects of stimuli, *F*(1,70) = 2.57, *p* = .114, arousal, *F*(1,70) = .00, *p* = .948, nor any interaction effects, *Fs*(1,70) < 1.03, *ps* > .315 (see Figure 8A and Table 6).

**Figure 8.**
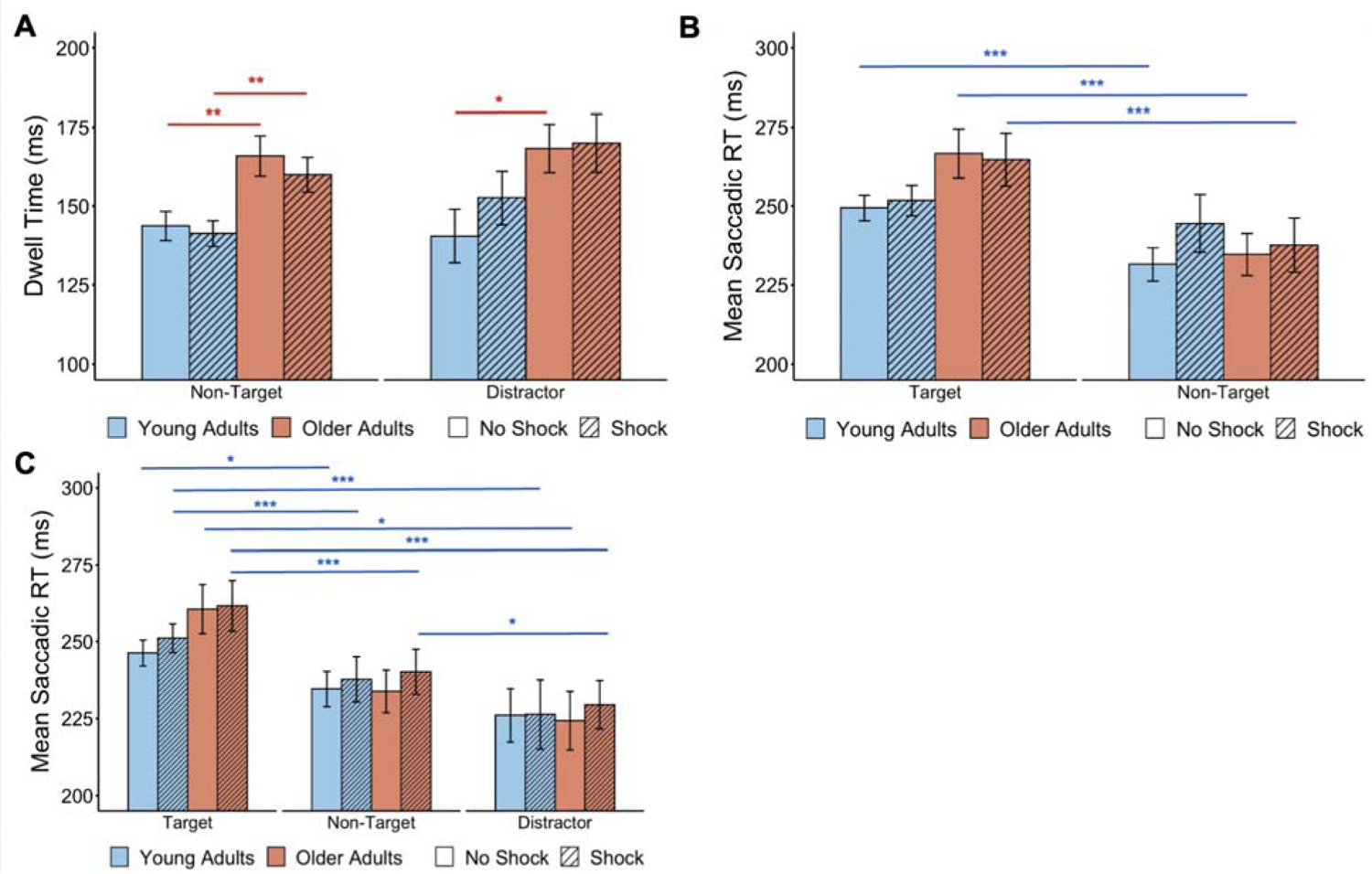
Threat of shock did not modulate dwell times or saccade latencies while in feature-search mode. Bar graphs depict (A) dwell time and mean saccadic reaction times in (B) distractor-absent and (C) distractor-present trials. Both age groups had longer fixation durations on the distractor compared to non-target shapes (see also Table 3). Both age groups showed longer mean saccade latencies when the direction of the saccade was the target (see blue asterisks). Error bars reflect standard error of the mean. \**p* < .05, \*\**p* < .01, \*\*\**p* < .001.

**Table 6.** Descriptive statistics and post-hoc t-test analyses during feature search for dependent measures dwell time and mean saccadic reaction times.

| Dwell time (ms) |  |  |  |  |  |
| --- | --- | --- | --- | --- | --- |
| Descriptive Statistics |  | No MANOVA interactions were significant |  |  |  |
| Young |  |  |  |  |  |
| M = 143.7, SE = 4.6 (NonTar, NS) |  |  |  |  |  |
| M = 141.3, SE = 4.1 (NonTar, S) |  |  |  |  |  |
| M = 140.5, SE = 8.5 (Dist, NS) |  |  |  |  |  |
| M = 152.6, SE = 8.4 (Dist, S) |  |  |  |  |  |
| Older |  |  |  |  |  |
| M = 165.9, SE = 6.4 (NonTar, NS) |  |  |  |  |  |
| M = 159.9, SE = 5.5 (NonTar, S) |  |  |  |  |  |
| M = 168.3, SE = 7.6 (Dist, NS) |  |  |  |  |  |
| M = 169.9, SE = 9.2 (Dist, S) |  |  |  |  |  |
| Mean saccadic reaction times on distractor-absent trials (ms) |  |  |  |  |  |
| Descriptive Statistics |  | Post-hoc t-test analyses | t | p | d |
| Young |  | Young vs. Older (Tar, NS) | 1.98 | .051 |  |
| M = 249.4, SE = 4.0 (Tar, NS) |  | Young vs. Older (Tar, S)* | 1.35 | .182 |  |
| M = 251.8, SE = 4.8 (Tar, S) |  | Young vs. Older (NonTar, NS) | .37 | .712 |  |
| M = 231.5, SE = 5.3 (NonTar, NS) |  | Young vs. Older (NonTar, S) | .75 | .459 |  |
| M = 244.5, SE = 9.1 (NonTar, S) |  |  |  |  |  |
| Older |  | Tar vs. NonTar (Young, NS)*** | 5.30 | <.001 | .597 |
| M = 266.7, SE = 7.7 (Tar, NS) |  | Tar vs. NonTar (Young, S) | .66 | .516 |  |
| M = 264.8, SE = 8.3 (Tar, S) |  | Tar vs. NonTar (Older, NS)*** | 9.89 | <.001 | .701 |
| M = 234.7, SE = 6.7 (NonTar, NS) |  | Tar vs. NonTar (Older, S)*** | 5.45 | <.001 | .534 |
| M = 237.6, SE = 8.6 (NonTar, S) |  |  |  |  |  |
|  |  | NS vs. S (YA, Tar) | .84 | .408 |  |
|  |  | NS vs. S (YA, NonTar) | 1.85 | .073 |  |
|  |  | NS vs. S (OA, Tar) | .61 | .545 |  |
|  |  | NS vs. S (OA, NonTar) | .64 | .526 |  |
| Mean saccadic reaction times on distractor-present trials (ms) |  |  |  |  |  |
| Descriptive Statistics |  | No MANOVA interactions were significant |  |  |  |
| Young |  |  |  |  |  |
| M = 246.3, SE = 4.2 (Tar, NS) |  |  |  |  |  |
| M = 251.1, SE = 4.7 (Tar, S) |  |  |  |  |  |
| M = 234.6, SE = 5.7 (NonTar, NS) |  |  |  |  |  |
| M = 237.8, SE = 7.4 (NonTar, S) |  |  |  |  |  |
| M = 226.0, SE = 8.7 (Dist, NS) |  |  |  |  |  |
| M = 226.3, SE = 11.3 (Dist, S) |  |  |  |  |  |
| Older |  |  |  |  |  |
| M = 260.6, SE = 8.0 (Tar, NS) |  |  |  |  |  |
| M = 261.6, SE = 8.3 (Tar, S) |  |  |  |  |  |
| M = 233.8, SE = 6.9 (NonTar, NS) |  |  |  |  |  |
| M = 240.2, SE = 7.3 (NonTar, S) |  |  |  |  |  |
| M = 224.3, SE = 9.5 (Dist, NS) |  |  |  |  |  |
| M = 229.5, SE = 7.9 (Dist, S) |  |  |  |  |  |

Note. M = mean, SE = standard error of the mean, Young = young adults, Older = Older adults, Abs = distractor-absent trials, Pres = distractor-present trials, Tar = target, NonTar = non-target, Dist = distractor, NS = no shock, S = shock. \*\*\*p < .001, \*\*p < .01, \*p < .05

Next, we investigated whether the threat of unpredictable shock modulated age differences in mean saccadic reaction times (sRT) on distractor-absent trials. A 2×2×2 MANOVA over factors age (young vs. older), stimuli (target vs. non-target), and arousal (no shock vs. shock) revealed a significant main effect of stimuli, *F*(1,70) = 52.26, *p* < .001, η*_p_^2^* = .427, but did not reveal a main effect of age, *F*(1,70) = .46, *p* = .500, nor arousal, *F*(1,70) = 2.40, *p* = .126 (see Figure 8B). However, we observed a significant age x stimuli interaction effect, *F*(1,70) = 10.40, *p* = .002, η*_p_^2^* = .129, and a significant arousal x stimuli interaction effect, *F*(1,70) = 4.66, *p* = .034, η*_p_^2^* = .062, but no other interaction effects, *Fs*(1,70) = 1.90, *p* > .172. Post-hoc analyses revealed that both young and older adults had higher mean sRTs when the first saccade was toward the target (see Table 6). Interestingly, there was marginal evidence that older adults took longer to initiate the first saccade when the direction was the target and that arousal delays mean sRTs in young adults when making a reflexive saccade toward a non-target stimulus (see Table 6).

Last, we conducted a 2×3×2 MANOVA over factors age (young vs. older), stimuli (target vs. non-target vs. distractor), and arousal (no shock vs. shock) on distractor-present trials. We observed a significant main effect of stimuli, *F*(2,140) = 20.07, *p* < .001, η*_p_^2^* = .251, but no main effects of age, *F*(1,70) = .38, *p* = .542, nor a main effect of arousal, *F*(1,70) = .28, *p* = .602 (see Figure 8C and Table 6). In addition, we did not observe any interaction effects, *Fs*(2,140) < .72, *ps* > .400.

### Threat of Unpredictable Shock on Pupillometry

Prior research have revealed the locus coeruleus-noradrenaline (LC-NE) is related to pupil size in humans (DiNuzzo et al., 2019; Grueschow et al., 2021, 2022; Murphy et al., 2014) and exhibited evidence for a causal link between locus coeruleus firing and pupil size in macaque monkeys (Joshi et al., 2016) and rodents (Breton-Provencher & Sur, 2019; Y. Liu et al., 2017). Furthermore, researchers have shown that evoked pupil dilations covary with transient increases in locus coeruleus activity (de Gee et al., 2017; Joshi et al., 2016; Joshi & Gold, 2022; Murphy et al., 2014; Varazzani et al., 2015). Thus, we investigated whether unpredictable threat of shock modulated both baseline pupil sizes and evoked pupil responses (pupil dilation responses) at the onset of the visual search array to evaluate whether changes in pupillometry characterized the tonic vs. phasic firing relationship of the LC-NE system. Given that participants were constantly under the threat of unpredictable shock throughout the entire run, emulating conditions of increased tonic noradrenergic activity through increased arousal, we hypothesized that the threat of shock would increase baseline pupil sizes and would thus decrease phasic noradrenergic responses to stimuli (as measured by evoked pupil dilations) following the Yerkes-Dodson curve (Aston-Jones & Cohen, 2005). We measured baseline resting pupil sizes within the 200 ms period before the onset of the visual search array, during which only the fixation cross was present on the screen. We defined pupil dilation responses as the peak change in pupil size from baseline resting states (%Change shows change from baseline pupil size). We averaged trials across all runs and then across all participants.

First, we investigated whether the threat of unpredictable shock modulated pupillometry measures during singleton search (see Figure 9A). A 2×2 MANOVA analysis over factors age (young vs. older) and arousal (no shock vs. shock) with baseline pupil size as the dependent measure revealed a main effect of age, *F*(1,66) = 27.03, *p* < .001, η*_p_^2^* = .291, main effect of arousal, *F*(1,66) = 7.49, *p* = .008, η*_p_ ^2^* = .102, but no interaction effect, *F*(1,66) = 1.42, *p* = .238 (see Figure 9B). Furthermore, we investigated whether arousal-induced changes in evoked pupil responses showed the opposite pattern compared with the effects on resting pupil size. A 2×2×3 MANOVA analysis over factors age (young vs. older), arousal (no shock vs. shock), and stimuli (target vs. non-target vs. distractor) revealed only a significant main effect of arousal, *F*(1,66) = 5.46, *p* = .023, η*_p_^2^* = .076, but no main effect of age, *F*(1,66) = .03, *p* = .862, stimuli, *F*(2,132) = .90, *p* = .408, nor any interaction effects, *Fs*(2,132) < 1.28, *ps* = .280 (see Figure 9C).

**Figure 9.**
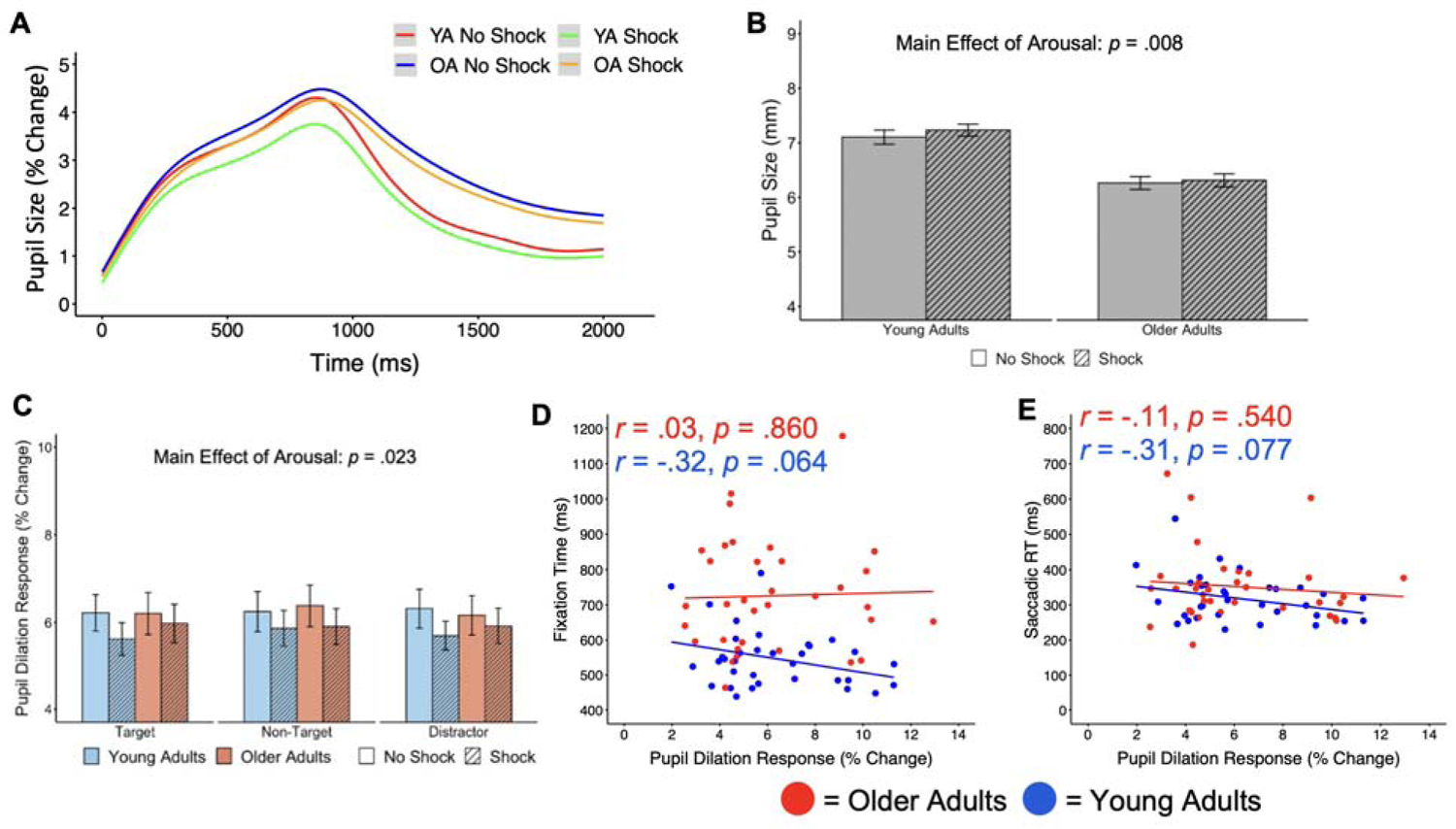
Threat of unpredictable shock increases resting pupil sizes and decreases evoked pupil responses during singleton search. (A) Starting from the onset of the visual search array during singleton search, the time series depicts changes in pupil size from baseline. Threat of unpredictable shock (B) increased resting pupil sizes and (C) decreased evoked pupil responses. In addition, (D-E) measures of oculomotor function exhibited a trending negative relationship with evoked pupil responses in young adults, but not in older adults.

Last, we investigated the relationship between measures of oculomotor function and evoked pupil responses, as both measures are characterized as functional indicators of the locus coeruleus-noradrenaline system (Joshi, 2023). We focused on fixation time as an overall measure of behavioral performance underlying eye movements and mean saccadic latencies as the primary locus coeruleus-related measures of oculomotor function (Joshi, 2023; Yamagishi & Furukawa, 2020). Interestingly, we observed that both measures of oculomotor function exhibited a marginally positive relationship with pupil dilation responses in young adults (see Figure 9D-E), that is phasic noradrenergic activity as seen in evoked pupil dilation responses exhibited a trending relationship with oculomotor function. We evaluated if there was a significant difference in correlation coefficients between the two age groups using the cocor R package (Diedenhofen & Musch, 2015). We observed no age differences in the dependent measure fixation time, *z* = 1.424, *p =* .144 (see Figure 9D), nor in saccadic reaction times, *z* = .827, *p =* .408 (see Figure 9E).

Next, we investigated whether the threat of unpredictable shock modulated pupillometry measures during feature search (see Figure 10A). A 2×2 MANOVA analysis over factors age (young vs. older) and arousal (no shock vs. shock) revealed a main effect of age, *F*(1,70) = 4.10, *p* = .047, η*_p_^2^* = .055, but did not reveal a main effect of arousal, *F*(1,70) = 1.58, *p* = .213, nor an interaction effect, *F*(1,70) = .06, *p* = .806 (see Figure 10B). A 2×2×3 MANOVA analysis over factors age (young vs. older), arousal (no shock vs. shock), and stimuli (target vs. non-target vs. distractor) over pupil dilation responses revealed a significant main effect of age, *F*(1,62) = 7.66, *p* = .007, η*_p_^2^* = .110, stimuli, *F*(2,124) = 5.58, *p* = .005, η*_p_^2^* = .083, but no main effect of arousal, *F*(1,62) = 1.78, *p* = .187 (see Figure 10C). In addition, we observed a significant age x stimuli interaction effect, *F*(2,124) = 6.31, *p* = .002, η*_p_^2^* = .092, with younger adults exhibiting significantly greater pupil dilation responses when the direction of the first saccade is toward the distractor. As in the singleton-search task, we explored whether measures of oculomotor function correlated with evoked pupil responses. We did not observe significant relationships in both age groups when engaging in feature search and observed no age differences between the correlation coefficients, *zs < .*615, *ps* > .539 (see Figure 10D-E).

**Figure 10.**
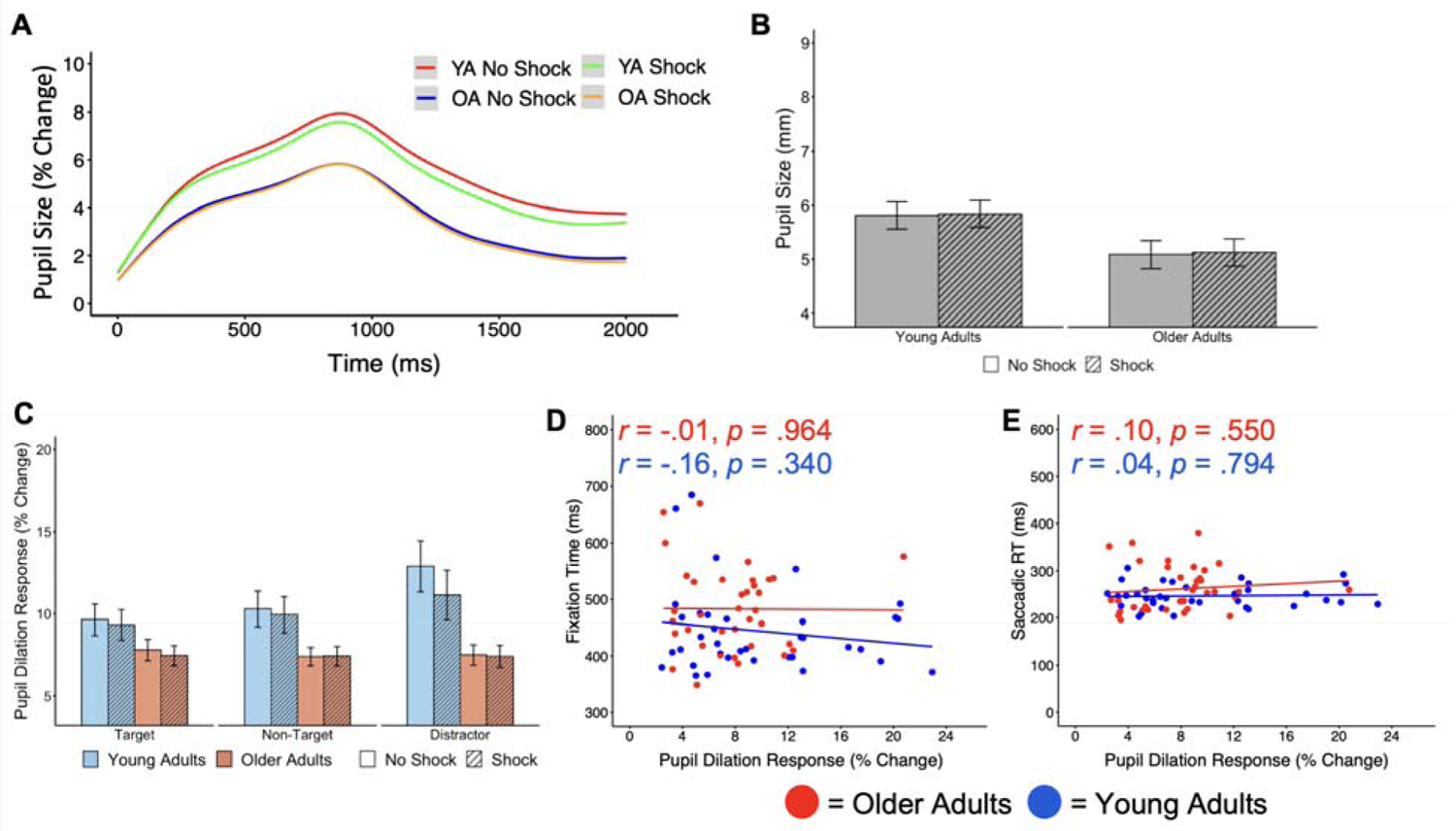
Threat of unpredictable shock does not modulate pupillometry metrics during feature search. (A) From the onset of the visual search array during feature search, we depict the time series of pupil size changes from baseline. Inducing arousal did not modulate (B) resting pupil sizes nor (C) pupil dilation responses. Furthermore, (D-E) measures of oculomotor function did not exhibit relationships with evoked pupil responses in both age groups.

### Sex Differences in Evoked Pupil Responses during Aging

Based on the tonic (baseline) noradrenergic hyperactivity hypothesis in aging, we hypothesized that evoked pupil dilation responses (phasic noradrenergic firing) would decrease in aging based on their inverse U-shaped relationship (Aston-Jones & Cohen, 2005). In both singleton-search and feature-search experiments, evoked pupil responses did not show significant correlations with age, *r =* −.03*, p* = .885, and *r =* −.02*, p* = .903, respectively. However, when plotting the relationship between evoked pupil responses and age mediated by sex, we observed opposite relationships in both experiments (see Figure 11A-B). Despite our experiments not designed to investigate sex differences, as seen in the uneven sample distribution of participants when separated by sex, we preliminarily investigated whether biological sex was a mediating variable on the effect of age on pupil dilation responses. We used the PROCESS macro for SPSS to regress age and gender on pupil dilation responses and found that the interaction of age and gender was trending but not significant during singleton search, *F*(1,30) = 2.44, *p* = .129, and during feature search, *F*(1,32) = 1.24, *p* = .274.

**Figure 11.**
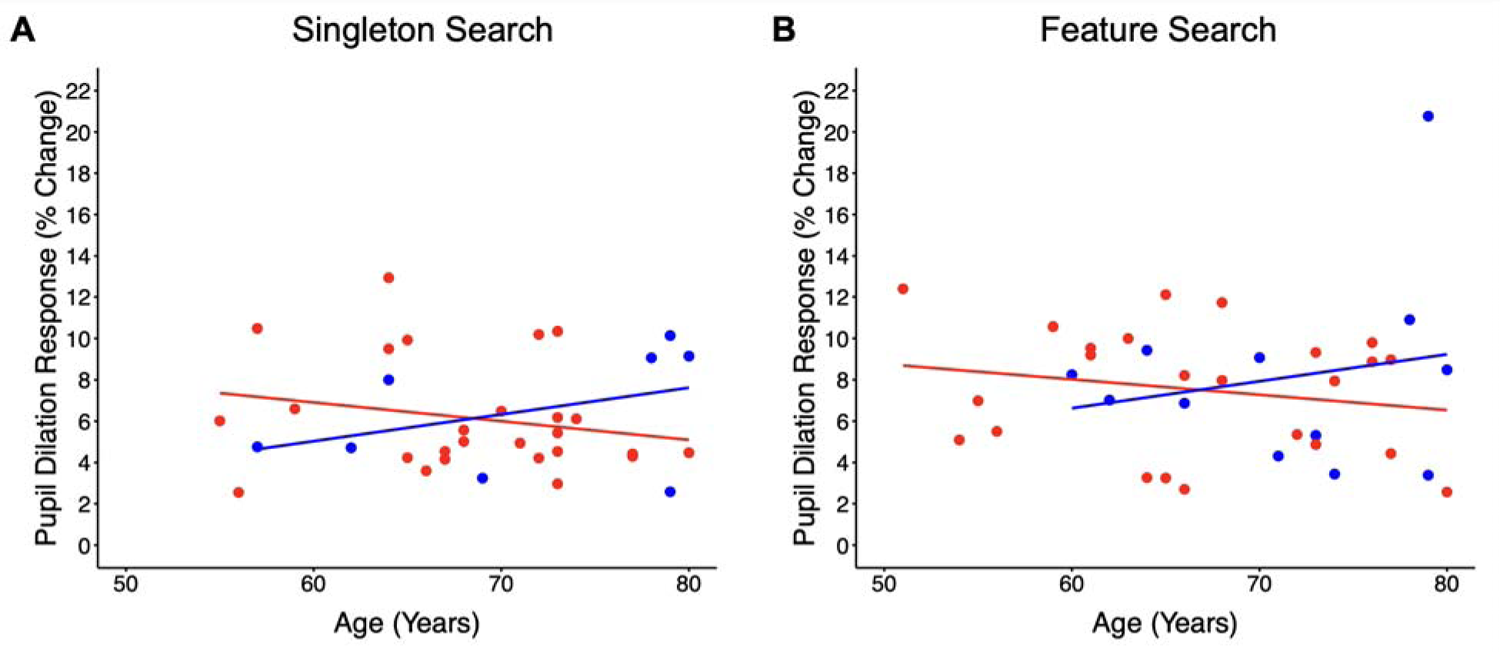
We did not observe a significant mediating effect of sex on the relationship between evoked pupil responses and age. We identified that evoked pupil responses did not relate to age during (A) singleton search and (B) feature search. However, when plotting relationships split by sex, we observed that the relationship showed opposite patterns. Given that our experiments were not powered to observe sex differences, the interaction of age and gender was also nonsignificant, but our data suggest the importance of investigating sex differences when assessing locus coeruleus function in aging. Red circles = female, blue circles = male.

## Discussion

We employed a systems neuroscience approach to investigate age differences in interactions between the locus coeruleus-noradrenaline (LC-NE) system and multiple attentional control networks (top-down vs. bottom-up; proactive vs. reactive), using pupillometry and oculomotor measures. Investigating eye movements were ideal for this purpose given its high reliability in both young and older adults (Kim, Grégoire, et al., 2024), the absence of confounding factors on aging compared with motor response times (Hollingworth & Bahle, 2020), and the known networks of interaction between the LC-NE system and oculomotor system in visual perception (Grefkes et al., 2010; Joshi, 2023; Kalwani et al., 2014; M. Lee et al., 2020; Sawaguchi & Kikuchi, 1998). Our primary goal was to investigate the mechanisms by which arousal differentially modulates attentional control in young and older adults. One of our primary research questions was to determine if arousal differentially modulates attention capture by the salient distractor in older adults compared with in young adults. Our MANOVA analysis revealed a marginal age by arousal interaction effect on first-saccade destinations to the salient distractor. Post-hoc t-tests analysis revealed that arousal significantly increases attention capture by salient stimuli in young adults, as previously observed using the identical unpredictable threat of shock manipulation (Kim & Anderson, 2020b). In older adults on the other hand, Bayesian statistics revealed moderate evidence in support of the null hypothesis, in that attention capture by salient stimuli is not modulated by arousal in older adults. Although we have replicated the arousal-biased competition effect of increased first saccades to the salient distractor in young adults, higher powered studies are needed to replicate our findings that arousal does not affect attention capture by the salient distractor in older adults. However, the observation that arousal does not modulate attentional priority for salient stimuli in older adults as seen with young adults is consistent with previous findings in the aging literature (Allen et al., 2017; Durbin et al., 2018; Gallant et al., 2022; T.-H. Lee et al., 2018; Svärd et al., 2014). By observing eye movements, we further revealed that arousal did not modulate the frequency of first saccades to any possible stimuli within the visual search array in older adults. Thus, in older adults, the lack of selectivity to salient stimuli when compared with non-salient stimuli under increased arousal is not due to equal changes in priority for multiple stimuli as seen in T.-H. Lee et al. (2018), but rather due to no changes in selectivity as in the neural mechanisms observed in Gallant et al. (2022). Furthermore, in young adults, arousal increases first saccade destinations to only the salient stimuli, demonstrating that the mechanism of arousal-biased competition in young adults is not a general shift from goal-oriented/top-down processing to reflexive/bottom-up processing, supporting the arousal-biased competition theory (Mather & Sutherland, 2011). Our findings indicate that aging may be changing or impeding how neural networks of arousal interact with bottom-up priority maps of saliency in the visual cortex that should be explored in future neuroimaging or electroencephalography studies.

Additionally, we observed that arousal did not modulate attention processing speeds with respect to the initiation of eye movements, reorienting, nor in overall fixation times. In the visual attention literature, improved performance under conditions of elevated arousal has commonly been determined by observing improvements in cognitive processing speed. In spatial cuing tasks that require orienting, stimuli that evoked higher emotional arousal led to faster responses in valid trials and also slower responses in invalid trials (T.-H. Lee et al., 2014; Sawada & Sato, 2015; Sutton & Lutz, 2019). When participants are presented with task-irrelevant arousing stimuli prior to the task, they can identify the target faster (T.-H. Lee et al., 2012; Sutherland & Mather, 2018; Zsido et al., 2020). Furthermore, dwell time is also longer for high arousal pictures with the length of dwell time being related to changes in pupil dilations (Astudillo et al., 2018). Interestingly, two studies have also revealed no behavioral effects of increased arousal despite showing significant changes in the late positive potential component of event-related potentials that indicate an electrophysiological arousal response (Ásgeirsson & Nieuwenhuis, 2017, 2019). These conflicting findings in the literature in addition to our divergent results from two unique search paradigms emphasize the importance of understanding why arousal does or does not modulate attention processing in specific task conditions and understanding the mechanisms underlying interactions between networks of arousal and each cognitive process. Our findings reveal that arousal does not speed neural processing nor slow mechanisms of attentional control by taking away cognitive resources. We argue that faster responses under arousal in the literature are solely because of changes in attentional priority of stimuli, naturally leading to faster orienting given task-specific demands such as in spatial orientation. Furthermore, our results suggest that arousal modulates priority maps in the visual cortex and increases attentional priority of salient stimuli as seen with the higher likelihood of fixating on the distractor during singleton search in the arousing than in the non-arousing blocks. The lack of changes in visual attention priority maps in older adults may be because of declines in functional connectivity between the locus coeruleus and the salience network (T.-H. Lee et al., 2020). However, steady state visual evoked potential studies show equal distractor suppression in the visual cortex when comparing young and older adults (Quigley et al., 2010; Quigley & Müller, 2014), suggesting that the age difference observed in this current study may be primarily driven by the interactions between the locus coeruleus-noradrenaline system and the visual cortex and not because of age-related declines in visual processing.

As in Ásgeirsson & Nieuwenhuis (2017, 2019), we also observed that arousal had null effects when completing visual search while engaged in feature-search mode (Bacon & Egeth, 1994). There are multiple possibilities for this repeated observation of a null effect of arousal-biased competition, even in young adults. First, Ásgeirsson & Nieuwenhuis (2019) suggest the plausibility that participants may be less aroused by certain experimental manipulations or environmental conditions. We observed differences in the effects of arousal on resting state pupil sizes and evoked pupil responses across experiments, with measures of pupillometry being non-significantly affected by arousal during feature search potentially explaining the lack of oculomotor effects. While it is possible that the sample populations we recruited for each experiment had general group differences in noradrenaline activity, we believe it is more likely that the task differences itself may have caused these differences in arousal-biased competition as observed with pupillometry. Resting state pupil sizes and evoked pupil dilation responses were significantly different across experiments, while they still maintained the same inverse pattern. Given that the singleton search task requires more mechanisms of cognitive control and is more difficult compared with feature search, we observed greater resting pupil sizes during singleton search and smaller evoked pupil dilation responses. Future studies would need to verify the hypothesis that increasing cognitive load or perhaps increasing the set size in visual search experiments will amplify or nullify the effect of arousal on attentional control. In addition, there was a key difference in how the attention system processed the salient distractor in the two experiments in our study. Our findings suggest that even when increasing attentional priority of salient stimuli and elevating the “attend-to-me” signal (Sawaki & Luck, 2010) through elevated arousal, top-down proactive suppression still overrides bottom-up attention capture (Chang & Egeth, 2021). It would be interesting to examine in future studies whether this increased attentional priority of the salient distractor under arousal further aided oculomotor suppression during feature search (e.g., increasing the set size of the visual search array to be able to observe greater suppression effects). Unfortunately, the neural networks controlling proactive vs. reactive mechanisms of attentional control are still debated (Geng, 2014; Hickey et al., 2009; Marini et al., 2016; Perri, 2020; van Belle et al., 2014; Vissers et al., 2016) and it is unclear whether the divergent effects are because the locus coeruleus-noradrenaline system differentially modulates potentially divergent neural networks. In our feature-search experiment, we also observed marginal effects of arousal on fixation times and a marginal arousal by first-saccade destination interaction in distractor-absent trials. It is plausible that arousal differentially modulates the speed of attention processing with delays in proactive mechanisms of attentional control, however, this is yet unclear with our data. Last, as also seen in Ásgeirsson & Nieuwenhuis (2017, 2019), our findings challenge the notion that arousal impairs top-down prioritization as observed in some prior findings (Gallant et al., 2022; Mather et al., 2016b; Sutherland et al., 2017). Even in distractor-absent trials in which goal-directed attentional control is not competing with bottom-up processes for saccade direction, arousal shows to have no effect on impairing goal-directed attentional control. Overall, the findings in both experiments both support and challenge multiple aspects of the arousal-biased competition theory (Mather & Sutherland, 2011) and also calls for higher-powered, future experiments to explore the neural networks and mechanisms by which arousal modulates attentional control.

A unique aspect of our experiments that contrasts with prior research investigating arousal-biased competition is the mode of arousal induction. Commonly in the fear and anxiety literature, researchers use the threat of unpredictable shock to emulate brain states of increased state anxiety (Balderston et al., 2017; Grillon et al., 2017; Robinson et al., 2013). Here, we used the threat of unpredictable shock to emulate states of tonic noradrenergic hyperactivity that is hypothesized to occur in older adults (Elman et al., 2017; Gannon & Wang, 2019; Mather, 2020; Weinshenker, 2018), given that the threat of unpredictable shock has shown to reduce alpha oscillations in the parietal lobe that mimics age-related changes (Balderston et al., 2017).

Although our oculomotor findings using unpredictable threat are consistent with prior behavioral results that induced arousal immediately prior to task presentation, there may be potential differences in the mechanisms by which threat of unpredictable shock vs. increased arousal prior to task presentation may act on cognition. The GANE model proposes that local noradrenaline hot spots occur in locations of prioritized representations to amplify pertinent processing (Mather et al., 2016b). However, are these hot spots sustained if the arousal is constant as in unpredictable threat? Do these hot spots disappear and only activate when an arousal-inducing stimulus is presented? If so, how long do they remain active? Further studies need to compare whether there are mechanistic differences in how unpredictable shock modulates arousal compared to momentary increases in arousal throughout a task. In addition, our findings both support and question the noradrenergic hyperactivity hypothesis in aging. Our oculomotor findings support the theory that dysregulated noradrenergic activity in aging leads to hyperactivity and not hypoactivity given that threat of unpredictable shock shifted attentional priority of salient stimuli in young adults but had no effect in older adults. If older adults exhibited hypoactivity, increases in arousal would significantly boost performance as predicted by the Yerkes-Dodson curve (Aston-Jones & Cohen, 2005). However, both young and older adults exhibited similar patterns of the effects of arousal on measures of pupillometry and we observed no age x arousal interaction effects. We hypothesize that although they exhibit noradrenergic hyperactivity, older adults are not so far on the extreme end to exhibit no changes in pupillometry due to increased arousal while this arousal-induced change from threat of unpredictable shock suffices to impair oculomotor performance. However, an important point to consider is the validity of evoked pupil responses as a measure of phasic noradrenergic activity in older adults. Our data show that young adults have greater pupil dilation responses to the distractor during feature search as expected, but not in older adults. Furthermore, we identified that two proposed measures of locus coeruleus function (oculomotor function and evoked pupil responses; Joshi, 2023) are marginally related in young adults but not in older adults. It is pertinent to evaluate the appropriateness of utilizing evoked pupil responses in older adults as a measure of phasic noradrenergic function in older adults, given what we know about dysregulated locus coeruleus function and decreased functional connectivity of the locus coeruleus in older adults.

Our collective findings inform and challenge many theories in the literature regarding the locus coeruleus-noradrenaline system, aging, and attentional control systems. In addition, our results challenge the aging, locus coeruleus-noradrenaline system, and neurodegenerative disease fields in two important ways. First, we observe a trend for opposing relationships of locus coeruleus function as seen with evoked pupil dilations and age when separating by sex. The locus coeruleus contains significantly more neurons in women compared with in men and estrogen has shown to increase tonic noradrenergic firing rates, potentially because of high sensitivity for corticotropin-releasing factor in locus coeruleus neurons (Bangasser et al., 2016; Curtis et al., 2006; Luckey et al., 2021). In addition, arousal has shown to have diverging effects in females and males (Sawada et al., 2014; Ycaza Herrera et al., 2019). Our preliminary findings further emphasize the importance of evaluating sex differences in locus coeruleus research. Second, eye movements are emerging as a potential measure of locus coeruleus function in humans (Joshi, 2023). Given the high reliability of oculomotor function even in older adults (Kim, Grégoire, et al., 2024), the establishment and validation of eye movements as a measure of locus coeruleus function may be critically useful to investigate changes in the locus coeruleus-noradrenaline system in aging and disease. Eye movements are effective in distinguishing between neurodegenerative diseases (Garbutt et al., 2008; Lage et al., 2020; Z. Liu et al., 2021) and also between patients with AD and prodromal stages such as mild cognitive impairment (Chehrehnegar et al., 2019, 2022; Opwonya et al., 2022; Peltsch et al., 2014; Wilcockson et al., 2019). Our results show the utility for eye movements to be a functional measure that may capture the progression of neurodegenerative diseases and predict the likelihood of transition to prodromal stages of dementia such as mild cognitive impairment.

## References

Allen, P. A., Lien, M.-C., & Jardin, E. (2017). Age-related emotional bias in processing two emotionally valenced tasks. Psychological Research, 81(1), 289–308. 10.1007/s00426-015-0711-8

Anderson, B. A., Kim, H., Kim, A. J., Liao, M.-R., Mrkonja, L., Clement, A., & Grégoire, L. (2021). The past, present, and future of selection history. Neuroscience & Biobehavioral Reviews, 130, 326–350. 10.1016/j.neubiorev.2021.09.004

Armstrong, K. M., Fitzgerald, J. K., & Moore, T. (2006). Changes in Visual Receptive Fields with Microstimulation of Frontal Cortex. Neuron, 50(5), 791–798. 10.1016/j.neuron.2006.05.010

Ásgeirsson, Á. G., & Nieuwenhuis, S. (2017). No arousal-biased competition in focused visuospatial attention. Cognition, 168, 191–204. 10.1016/j.cognition.2017.07.001

Ásgeirsson, Á. G., & Nieuwenhuis, S. (2019). Effects of arousal on biased competition in attention and short-term memory. Attention, Perception, & Psychophysics, 81(6), 1901– 1912. 10.3758/s13414-019-01756-x

Aston-Jones, G., & Cohen, J. D. (2005). AN INTEGRATIVE THEORY OF LOCUS COERULEUS-NOREPINEPHRINE FUNCTION: Adaptive Gain and Optimal Performance. Annual Review of Neuroscience, 28(1), 403–450. 10.1146/annurev.neuro.28.061604.135709

Aston-Jones, G., Rajkowski, J., & Cohen, J. (1999). Role of locus coeruleus in attention and behavioral flexibility. Biological Psychiatry, 46(9), 1309–1320. 10.1016/s0006-3223(99)00140-7

Aston-Jones, G., Rajkowski, J., & Cohen, J. (2000). Locus coeruleus and regulation of behavioral flexibility and attention. In Progress in Brain Research (Vol. 126, pp. 165–182). Elsevier. 10.1016/S0079-6123(00)26013-5

Astudillo, C., Muñoz, K., & Maldonado, P. E. (2018). Emotional Content Modulates Attentional Visual Orientation During Free Viewing of Natural Images. Frontiers in Human Neuroscience, 12. 10.3389/fnhum.2018.00459

Bacon, W. F., & Egeth, H. E. (1994). Overriding stimulus-driven attentional capture. Perception & Psychophysics, 55(5), 485–496. 10.3758/BF03205306

Balderston, N. L., Hale, E., Hsiung, A., Torrisi, S., Holroyd, T., Carver, F. W., Coppola, R., Ernst, M., & Grillon, C. (2017). Threat of shock increases excitability and connectivity of the intraparietal sulcus. eLife, 6, e23608. 10.7554/eLife.23608

Bangasser, D. A., Wiersielis, K. R., & Khantsis, S. (2016). Sex Differences in the Locus Coeruleus-Norepinephrine System and its Regulation by Stress. Brain Research, 1641(Pt B), 177–188. 10.1016/j.brainres.2015.11.021

Berridge, C. W. (2008). Noradrenergic modulation of arousal. Brain Research Reviews, 58(1), 1– 17. 10.1016/j.brainresrev.2007.10.013

Berridge, C. W., & Waterhouse, B. D. (2003). The locus coeruleus-noradrenergic system: Modulation of behavioral state and state-dependent cognitive processes. Brain Research. Brain Research Reviews, 42(1), 33–84. 10.1016/s0165-0173(03)00143-7

Bouret, S., & Sara, S. J. (2004). Reward expectation, orientation of attention and locus coeruleus-medial frontal cortex interplay during learning. The European Journal of Neuroscience, 20(3), 791–802. 10.1111/j.1460-9568.2004.03526.x

Brainard, D. H. (1997). The Psychophysics Toolbox. Spatial Vision, 10(4), 433–436. 10.1163/156856897X00357

Breton-Provencher, V., & Sur, M. (2019). Active control of arousal by a locus coeruleus GABAergic circuit. Nature Neuroscience, 22(2), 218–228. 10.1038/s41593-018-0305-z

Bruce, C. J., & Goldberg, M. E. (1985). Primate frontal eye fields. I. Single neurons discharging before saccades. Journal of Neurophysiology, 53(3), 603–635. 10.1152/jn.1985.53.3.603

Campbell, K. L., Grady, C. L., Ng, C., & Hasher, L. (2012). Age differences in the frontoparietal cognitive control network: Implications for distractibility. Neuropsychologia, 50(9), 2212– 2223.

Case, G. R., & Ferrera, V. P. (2007). Coordination of smooth pursuit and saccade target selection in monkeys. Journal of Neurophysiology, 98(4), 2206–2214. 10.1152/jn.00021.2007

Cassidy, C. M., Therriault, J., Pascoal, T. A., Cheung, V., Savard, M., Tuominen, L., Chamoun, M., McCall, A., Celebi, S., Lussier, F., Massarweh, G., Soucy, J.-P., Weinshenker, D., Tardif, C., Ismail, Z., Gauthier, S., & Rosa-Neto, P. (2022). Association of locus coeruleus integrity with Braak stage and neuropsychiatric symptom severity in Alzheimer’s disease. Neuropsychopharmacology, 47(5), Article 5. 10.1038/s41386-022-01293-6

Chang, S., & Egeth, H. E. (2021). Can salient stimuli really be suppressed? Attention, Perception, & Psychophysics, 83(1), Article 1. 10.3758/s13414-020-02207-8

Chehrehnegar, N., Nejati, V., Shati, M., Esmaeili, M., Rezvani, Z., Haghi, M., & Foroughan, M. (2019). Behavioral and cognitive markers of mild cognitive impairment: Diagnostic value of saccadic eye movements and Simon task. Aging Clinical and Experimental Research, 31(11), 1591–1600. 10.1007/s40520-019-01121-w

Chehrehnegar, N., Shati, M., Esmaeili, M., & Foroughan, M. (2022). Executive function deficits in mild cognitive impairment: Evidence from saccade tasks. Aging & Mental Health, 26(5), 1001–1009. 10.1080/13607863.2021.1913471

Clewett, D. V., Huang, R., Velasco, R., Lee, T.-H., & Mather, M. (2018). Locus Coeruleus Activity Strengthens Prioritized Memories Under Arousal. Journal of Neuroscience, 38(6), 1558–1574. 10.1523/JNEUROSCI.2097-17.2017

Clewett, D. V., Lee, T.-H., Greening, S., Ponzio, A., Margalit, E., & Mather, M. (2016). Neuromelanin marks the spot: Identifying a locus coeruleus biomarker of cognitive reserve in healthy aging. Neurobiology of Aging, 37, 117–126. 10.1016/j.neurobiolaging.2015.09.019

Corbetta, M., Patel, G., & Shulman, G. L. (2008). The Reorienting System of the Human Brain: From Environment to Theory of Mind. Neuron, 58(3), 306–324. 10.1016/j.neuron.2008.04.017

Corbetta, M., & Shulman, G. L. (2002). Control of goal-directed and stimulus-driven attention in the brain. Nature Reviews Neuroscience, 3(3), 201–215. 10.1038/nrn755

Curtis, A. L., Bethea, T., & Valentino, R. J. (2006). Sexually Dimorphic Responses of the Brain Norepinephrine System to Stress and Corticotropin-Releasing Factor. Neuropsychopharmacology, 31(3), 544–554. 10.1038/sj.npp.1300875

Dahl, M. J., Bachman, S. L., Dutt, S., Düzel, S., Bodammer, N. C., Lindenberger, U., Kühn, S., Werkle-Bergner, M., & Mather, M. (2023). The integrity of dopaminergic and noradrenergic brain regions is associated with different aspects of late-life memory performance. Nature Aging, 1–16. 10.1038/s43587-023-00469-z

Dahl, M. J., Mather, M., Düzel, S., Bodammer, N. C., Lindenberger, U., Kühn, S., & Werkle-Bergner, M. (2019). Rostral locus coeruleus integrity is associated with better memory performance in older adults. Nature Human Behaviour, 3(11), Article 11. 10.1038/s41562-019-0715-2

Dahl, M. J., Mather, M., Werkle-Bergner, M., Kennedy, B. L., Guzman, S., Hurth, K., Miller, C. A., Qiao, Y., Shi, Y., Chui, H. C., & Ringman, J. M. (2022). Locus coeruleus integrity is related to tau burden and memory loss in autosomal-dominant Alzheimer’s disease. Neurobiology of Aging, 112, 39–54. 10.1016/j.neurobiolaging.2021.11.006

de Gee, J. W., Colizoli, O., Kloosterman, N. A., Knapen, T., Nieuwenhuis, S., & Donner, T. H. (2017). Dynamic modulation of decision biases by brainstem arousal systems. eLife, 6, e23232. 10.7554/eLife.23232

Desimone, R., & Duncan, J. (1995). Neural Mechanisms of Selective Visual Attention. Annual Review of Neuroscience, 18(1), 193–222. 10.1146/annurev.ne.18.030195.001205

Diedenhofen, B., & Musch, J. (2015). cocor: A Comprehensive Solution for the Statistical Comparison of Correlations. PLOS ONE, 10(4), e0121945. 10.1371/journal.pone.0121945

DiNuzzo, M., Mascali, D., Moraschi, M., Bussu, G., Maugeri, L., Mangini, F., Fratini, M., & Giove, F. (2019). Brain Networks Underlying Eye’s Pupil Dynamics. Frontiers in Neuroscience, 13, 965. 10.3389/fnins.2019.00965

Durbin, K. A., Clewett, D., Huang, R., & Mather, M. (2018). Age differences in selective memory of goal-relevant stimuli under threat. Emotion, 18(6), 906–911. 10.1037/emo0000398

Elman, J. A., Panizzon, M. S., Hagler, D. J., Eyler, L. T., Granholm, E. L., Fennema-Notestine, C., Lyons, M. J., McEvoy, L. K., Franz, C. E., Dale, A. M., & Kremen, W. S. (2017). Task-evoked pupil dilation and BOLD variance as indicators of locus coeruleus dysfunction. Cortex, 97, 60– 69. 10.1016/j.cortex.2017.09.025

Elman, J. A., Puckett, O. K., Beck, A., Fennema-Notestine, C., Cross, L. K., Dale, A. M., Eglit, G. M. L., Eyler, L. T., Gillespie, N. A., Granholm, E. L., Gustavson, D. E., Hagler, D. J., Hatton, S. N., Hauger, R., Jak, A. J., Logue, M. W., McEvoy, L. K., McKenzie, R. E., Neale, M. C., … Kremen, W. S. (2021). MRI-assessed locus coeruleus integrity is heritable and associated with multiple cognitive domains, mild cognitive impairment, and daytime dysfunction. Alzheimer’s & Dementia: The Journal of the Alzheimer’s Association, 17(6), Article 6. 10.1002/alz.12261

Gallant, S. N., Kennedy, B. L., Bachman, S. L., Huang, R., Cho, C., Lee, T.-H., & Mather, M. (2022). Behavioral and fMRI evidence that arousal enhances bottom-up selectivity in young but not older adults. Neurobiology of Aging, 120, 149–166. 10.1016/j.neurobiolaging.2022.08.006

Gamboz, N., Zamarian, S., & Cavallero, C. (2010). Age-Related Differences in the Attention Network Test (ANT). Experimental Aging Research, 36(3), 287–305. 10.1080/0361073X.2010.484729

Gannon, M., & Wang, Q. (2019). Complex noradrenergic dysfunction in Alzheimer’s disease: Low norepinephrine input is not always to blame. Brain Research, 1702, 12–16. 10.1016/j.brainres.2018.01.001

Garbutt, S., Matlin, A., Hellmuth, J., Schenk, A. K., Johnson, J. K., Rosen, H., Dean, D., Kramer, J., Neuhaus, J., Miller, B. L., Lisberger, S. G., & Boxer, A. L. (2008). Oculomotor function in frontotemporal lobar degeneration, related disorders and Alzheimer’s disease. Brain, 131(5), 1268–1281. 10.1093/brain/awn047

Gaspelin, N., Leonard, C. J., & Luck, S. J. (2017). Suppression of overt attentional capture by salient-but-irrelevant color singletons. Attention, Perception, & Psychophysics, 79(1), 45–62. 10.3758/s13414-016-1209-1

Geerligs, L., Renken, R. J., Saliasi, E., Maurits, N. M., & Lorist, M. M. (2015). A Brain-Wide Study of Age-Related Changes in Functional Connectivity. Cerebral Cortex, 25(7), 1987–1999. 10.1093/cercor/bhu012

Geng, J. J. (2014). Attentional Mechanisms of Distractor Suppression. Current Directions in Psychological Science, 23(2), 147–153. 10.1177/0963721414525780

Ghosh, S., & Maunsell, J. H. R. (2022). Locus coeruleus norepinephrine selectively controls visual attention [Preprint]. Neuroscience. 10.1101/2022.09.30.510394

Grady, C., Sarraf, S., Saverino, C., & Campbell, K. (2016). Age differences in the functional interactions among the default, frontoparietal control, and dorsal attention networks. Neurobiology of Aging, 41, 159–172. 10.1016/j.neurobiolaging.2016.02.020

Grefkes, C., Wang, L. E., Eickhoff, S. B., & Fink, G. R. (2010). Noradrenergic modulation of cortical networks engaged in visuomotor processing. Cerebral Cortex (New York, N.Y.: 1991), 20(4), 783–797. 10.1093/cercor/bhp144

Grillon, C., O’Connell, K., Lieberman, L., Alvarez, G., Geraci, M., Pine, D. S., & Ernst, M. (2017). Distinct responses to predictable and unpredictable threat in anxiety pathologies: Effect of panic attack. Biological Psychiatry. Cognitive Neuroscience and Neuroimaging, 2(7), 575–581. 10.1016/j.bpsc.2016.08.005

Grueschow, M., Kleim, B., & Ruff, C. C. (2022). Functional Coupling of the Locus Coeruleus Is Linked to Successful Cognitive Control. Brain Sciences, 12(3), Article 3. 10.3390/brainsci12030305

Grueschow, M., Stenz, N., Thörn, H., Ehlert, U., Breckwoldt, J., Brodmann Maeder, M., Exadaktylos, A. K., Bingisser, R., Ruff, C. C., & Kleim, B. (2021). Real-world stress resilience is associated with the responsivity of the locus coeruleus. Nature Communications, 12(1), 2275. 10.1038/s41467-021-22509-1

Hämmerer, D., Callaghan, M. F., Hopkins, A., Kosciessa, J., Betts, M., Cardenas-Blanco, A., Kanowski, M., Weiskopf, N., Dayan, P., Dolan, R. J., & Düzel, E. (2018). Locus coeruleus integrity in old age is selectively related to memories linked with salient negative events. Proceedings of the National Academy of Sciences of the United States of America, 115(9), 2228–2233. 10.1073/pnas.1712268115

Hasher, L., & Zacks, R. T. (1988). Working Memory, Comprehension, and Aging: A Review and a New View. In G. H. Bower (Ed.), Psychology of Learning and Motivation (Vol. 22, pp. 193– 225). Academic Press. 10.1016/S0079-7421(08)60041-9

Hickey, C., Di Lollo, V., & McDonald, J. J. (2009). Electrophysiological Indices of Target and Distractor Processing in Visual Search. Journal of Cognitive Neuroscience, 21(4), 760–775. 10.1162/jocn.2009.21039

Hollingworth, A., & Bahle, B. (2020). Eye Tracking in Visual Search Experiments. In S. Pollmann (Ed.), Spatial Learning and Attention Guidance (pp. 23–35). Springer US. 10.1007/7657_2019_30

Howard, J. H. Jr., Howard, D. V., Dennis, N. A., Yankovich, H., & Vaidya, C. J. (2004). Implicit Spatial Contextual Learning in Healthy Aging. Neuropsychology, 18(1), 124–134. 10.1037/0894-4105.18.1.124

Itti, L., & Koch, C. (2000). A saliency-based search mechanism for overt and covert shifts of visual attention. Vision Research, 40(10), 1489–1506. 10.1016/S0042-6989(99)00163-7

Itti, L., & Koch, C. (2001). Computational modelling of visual attention. Nature Reviews. Neuroscience, 2(3), 194–203. 10.1038/35058500

Jacobs, H. I. L., Wiese, S., van de Ven, V., Gronenschild, E. H. B. M., Verhey, F. R. J., & Matthews, P. M. (2015). Relevance of parahippocampal-locus coeruleus connectivity to memory in early dementia. Neurobiology of Aging, 36(2), 618–626. 10.1016/j.neurobiolaging.2014.10.041

Jia, X., Gao, C., Wang, Y., Han, M., Cui, L., & Guo, C. (2020). Emotional arousal influences remembrance of goal-relevant stimuli. Emotion, 20(8), 1357–1368. 10.1037/emo0000657

Joshi, S. (2023). The impact of age and Alzheimer’s disease on locus coeruleus mediated neuromodulation of neural circuits and goal-directed behavior. 10.31234/osf.io/bqwte

Joshi, S., & Gold, J. I. (2022). Context-dependent relationships between locus coeruleus firing patterns and coordinated neural activity in the anterior cingulate cortex. Elife, 11, e63490.

Joshi, S., Li, Y., Kalwani, R. M., & Gold, J. I. (2016). Relationships between Pupil Diameter and Neuronal Activity in the Locus Coeruleus, Colliculi, and Cingulate Cortex. Neuron, 89(1), 221– 234. 10.1016/j.neuron.2015.11.028

Kalwani, R. M., Joshi, S., & Gold, J. I. (2014). Phasic Activation of Individual Neurons in the Locus Ceruleus/Subceruleus Complex of Monkeys Reflects Rewarded Decisions to Go But Not Stop. Journal of Neuroscience, 34(41), 13656–13669. 10.1523/JNEUROSCI.2566-14.2014

Kennedy, B. L., & Mather, M. (2019). Neural mechanisms underlying age-related changes in attentional selectivity. In The aging brain: Functional adaptation across adulthood (pp. 45– 72). American Psychological Association. 10.1037/0000143-003

Kim, A., & Anderson, B. (2020a). Arousal-Biased Competition Explains Reduced Distraction by Reward Cues under Threat. eNeuro, 7(4), ENEURO.0099-20.2020. 10.1523/ENEURO.0099-20.2020

Kim, A., & Anderson, B. (2020b). Threat reduces value-driven but not salience-driven attentional capture. Emotion, 20, 874–889. 10.1037/emo0000599

Kim, A., & Anderson, B. (2022). Systemic effects of selection history on learned ignoring. Psychonomic Bulletin & Review, 29(4), Article 4. 10.3758/s13423-021-02050-4

Kim, A., Grégoire, L., & Anderson, B. A. (2024). Reliably Measuring Learning-Dependent Distractor Suppression with Eye Tracking (p. 2024.02.23.581757). bioRxiv. 10.1101/2024.02.23.581757

Kim, A., Lee, D., & Anderson, B. (2021). The influence of threat on the efficiency of goal-directed attentional control. Psychological Research, 85(3), 980–986. 10.1007/s00426-020-01321-4

Kim, A., Senior, J., Chu, S., & Mather, M. (2024). Aging Impairs Reactive Attentional Control but Not Proactive Distractor Inhibition. Journal of Experimental Psychology: General.

Kubo-Kawai, N., & Kawai, N. (2010). Elimination of the enhanced Simon effect for older adults in a three-choice situation: Ageing and the Simon effect in a go/no-go Simon task. Quarterly Journal of Experimental Psychology, 63(3), 452–464. 10.1080/17470210902990829

Lage, C., López-García, S., Bejanin, A., Kazimierczak, M., Aracil-Bolaños, I., Calvo-Córdoba, A., Pozueta, A., García-Martínez, M., Fernández-Rodríguez, A., Bravo-González, M., Jiménez-Bonilla, J., Banzo, I., Irure-Ventura, J., Pegueroles, J., Illán-Gala, I., Fortea, J., Rodríguez-Rodríguez, E., Lleó-Bisa, A., García-Cena, C. E., & Sánchez-Juan, P. (2020). Distinctive Oculomotor Behaviors in Alzheimer’s Disease and Frontotemporal Dementia. Frontiers in Aging Neuroscience, 12. 10.3389/fnagi.2020.603790

Laretzaki, G., Plainis, S., Argyropoulos, S., Pallikaris, I., & Bitsios, P. (2010). Threat and anxiety affect visual contrast perception. Journal of Psychopharmacology, 24(5), 667–675. 10.1177/0269881108098823

Lee, M., Mueller, A., & Moore, T. (2020). Differences in Noradrenaline Receptor Expression Across Different Neuronal Subtypes in Macaque Frontal Eye Field. Frontiers in Neuroanatomy, 14. 10.3389/fnana.2020.574130

Lee, T.-H., Greening, S. G., Ueno, T., Clewett, D., Ponzio, A., Sakaki, M., & Mather, M. (2018). Arousal increases neural gain via the locus coeruleus–noradrenaline system in younger adults but not in older adults. Nature Human Behaviour, 2(5), Article 5. 10.1038/s41562-018-0344-1

Lee, T.-H., Itti, L., & Mather, M. (2012). Evidence for Arousal-Biased Competition in Perceptual Learning. Frontiers in Psychology, 3. https://www.frontiersin.org/articles/10.3389/fpsyg.2012.00241

Lee, T.-H., Kim, S. H., Katz, B., & Mather, M. (2020). The Decline in Intrinsic Connectivity Between the Salience Network and Locus Coeruleus in Older Adults: Implications for Distractibility. Frontiers in Aging Neuroscience, 12. https://www.frontiersin.org/articles/10.3389/fnagi.2020.00002

Lee, T.-H., Sakaki, M., Cheng, R., Velasco, R., & Mather, M. (2014). Emotional arousal amplifies the effects of biased competition in the brain. Social Cognitive and Affective Neuroscience, 9(12), 2067–2077. 10.1093/scan/nsu015

Lega, C., Di Caro, V., Strina, V., & Daini, R. (2023). Age-related differences in the statistical learning of target selection and distractor suppression. Psychology and Aging, 38(3), Article 3. 10.1037/pag0000735

Li, L., Feng, X., Zhou, Z., Zhang, H., Shi, Q., Lei, Z., Shen, P., Yang, Q., Zhao, B., Chen, S., Li, L., Zhang, Y., Wen, P., Lu, Z., Li, X., Xu, F., & Wang, L. (2018). Stress Accelerates Defensive Responses to Looming in Mice and Involves a Locus Coeruleus-Superior Colliculus Projection. Current Biology: CB, 28(6), 859–871.e5. 10.1016/j.cub.2018.02.005

Liu, K. Y., Kievit, R. A., Tsvetanov, K. A., Betts, M. J., Düzel, E., Rowe, J. B., Howard, R., & Hämmerer, D. (2020). Noradrenergic-dependent functions are associated with age-related locus coeruleus signal intensity differences. Nature Communications, 11(1), Article 1. 10.1038/s41467-020-15410-w

Liu, Y., Rodenkirch, C., Moskowitz, N., Schriver, B., & Wang, Q. (2017). Dynamic Lateralization of Pupil Dilation Evoked by Locus Coeruleus Activation Results from Sympathetic, Not Parasympathetic, Contributions. Cell Reports, 20(13), 3099–3112. 10.1016/j.celrep.2017.08.094

Liu, Z., Yang, Z., Gu, Y., Liu, H., & Wang, P. (2021). The effectiveness of eye tracking in the diagnosis of cognitive disorders: A systematic review and meta-analysis. PLOS ONE, 16(7), e0254059. 10.1371/journal.pone.0254059

Loughlin, S. E., Foote, S. L., & Grzanna, R. (1986). Efferent projections of nucleus locus coeruleus: Morphologic subpopulations have different efferent targets. Neuroscience, 18(2), 307–319. 10.1016/0306-4522(86)90156-9

Luckey, A. M., Robertson, I. H., Lawlor, B., Mohan, A., & Vanneste, S. (2021). Sex Differences in Locus Coeruleus: A Heuristic Approach That May Explain the Increased Risk of Alzheimer’s Disease in Females. Journal of Alzheimer’s Disease, 83(2), 505–522. 10.3233/JAD-210404

Lustig, C., Hasher, L., & Zacks, R. T. (2007). Inhibitory deficit theory: Recent developments in a “new view.” In Inhibition in cognition (pp. 145–162). American Psychological Association. 10.1037/11587-008

Lyon, J., Scialfa, C., Cordazzo, S., & Bubric, K. (2014). Contextual cuing: The effects of stimulus variation, intentionality, and aging. Canadian Journal of Experimental Psychology / Revue Canadienne de Psychologie Expérimentale, 68(2), 111–121. 10.1037/cep0000007

Madden, D. J. (2007). Aging and Visual Attention. Current Directions in Psychological Science, 16(2), 70–74. 10.1111/j.1467-8721.2007.00478.x

Mahoney, J. R., Verghese, J., Goldin, Y., Lipton, R., & Holtzer, R. (2010). Alerting, orienting, and executive attention in older adults. Journal of the International Neuropsychological Society, 16(5), 877–889. 10.1017/S1355617710000767

Maness, E. B., Burk, J. A., McKenna, J. T., Schiffino, F. L., Strecker, R. E., & McCoy, J. G. (2022). Role of the locus coeruleus and basal forebrain in arousal and attention. Brain Research Bulletin, 188, 47–58. 10.1016/j.brainresbull.2022.07.014

Marini, F., Demeter, E., Roberts, K. C., Chelazzi, L., & Woldorff, M. G. (2016). Orchestrating Proactive and Reactive Mechanisms for Filtering Distracting Information: Brain-Behavior Relationships Revealed by a Mixed-Design fMRI Study. The Journal of Neuroscience, 36(3), 988–1000. 10.1523/JNEUROSCI.2966-15.2016

Mather, M. (2020). How Arousal-Related Neurotransmitter Systems Compensate for Age-Related Decline. In A. Gutchess & A. K. Thomas (Eds.), The Cambridge Handbook of Cognitive Aging: A Life Course Perspective (pp. 101–120). Cambridge University Press. 10.1017/9781108552684.007

Mather, M., Clewett, D., Sakaki, M., & Harley, C. W. (2016a). GANEing traction: The broad applicability of NE hotspots to diverse cognitive and arousal phenomena. The Behavioral and Brain Sciences, 39, e228. 10.1017/S0140525X16000017

Mather, M., Clewett, D., Sakaki, M., & Harley, C. W. (2016b). Norepinephrine ignites local hotspots of neuronal excitation: How arousal amplifies selectivity in perception and memory. Behavioral and Brain Sciences, 39, e200. 10.1017/S0140525X15000667

Mather, M., & Harley, C. W. (2016). The Locus Coeruleus: Essential for Maintaining Cognitive Function and the Aging Brain. Trends in Cognitive Sciences, 20(3), 214–226. 10.1016/j.tics.2016.01.001

Mather, M., & Sutherland, M. R. (2011). Arousal-Biased Competition in Perception and Memory. Perspectives on Psychological Science, 6(2), 114–133. 10.1177/1745691611400234

McBurney-Lin, J., Lu, J., Zuo, Y., & Yang, H. (2019). Locus Coeruleus-Norepinephrine Modulation of Sensory Processing and Perception: A Focused Review. Neuroscience and Biobehavioral Reviews, 105, 190–199. 10.1016/j.neubiorev.2019.06.009

McDonough, I. M., Wood, M. M., & Miller, Jr., W. S. (2019). A Review on the Trajectory of Attentional Mechanisms in Aging and the Alzheimer’s Disease Continuum through the Attention Network Test. The Yale Journal of Biology and Medicine, 92(1), 37–51.

Murphy, P. R., O’Connell, R. G., O’Sullivan, M., Robertson, I. H., & Balsters, J. H. (2014). Pupil diameter covaries with BOLD activity in human locus coeruleus. Human Brain Mapping, 35(8), 4140–4154. 10.1002/hbm.22466

Nashiro, K., Sakaki, M., Braskie, M. N., & Mather, M. (2017). Resting-state networks associated with cognitive processing show more age-related decline than those associated with emotional processing. Neurobiology of Aging, 54, 152–162. 10.1016/j.neurobiolaging.2017.03.003

Nummela, S. U., & Krauzlis, R. J. (2011). Superior Colliculus Inactivation Alters the Weighted Integration of Visual Stimuli. The Journal of Neuroscience, 31(22), 8059–8066. 10.1523/JNEUROSCI.5480-10.2011

Opwonya, J., Doan, D. N. T., Kim, S. G., Kim, J. I., Ku, B., Kim, S., Park, S., & Kim, J. U. (2022). Saccadic Eye Movement in Mild Cognitive Impairment and Alzheimer’s Disease: A Systematic Review and Meta-Analysis. Neuropsychology Review, 32(2), 193–227. 10.1007/s11065-021-09495-3

Padmala, S., & Pessoa, L. (2008). Affective Learning Enhances Visual Detection and Responses in Primary Visual Cortex. The Journal of Neuroscience, 28(24), 6202–6210. 10.1523/JNEUROSCI.1233-08.2008

Peltsch, A., Hemraj, A., Garcia, A., & Munoz, D. P. (2014). Saccade deficits in amnestic mild cognitive impairment resemble mild Alzheimer’s disease. European Journal of Neuroscience, 39(11), 2000–2013. 10.1111/ejn.12617

Perri, R. L. (2020). Is there a proactive and a reactive mechanism of inhibition? Towards an executive account of the attentional inhibitory control model. Behavioural Brain Research, 377, 112243. 10.1016/j.bbr.2019.112243

Phelps, E. A., Ling, S., & Carrasco, M. (2006). Emotion facilitates perception and potentiates the perceptual benefits of attention. Psychological Science, 17(4), 292–299. 10.1111/j.1467-9280.2006.01701.x

Pickel, V. M., Segal, M., & Bloom, F. E. (1974). A radioautographic study of the efferent pathways of the nucleus locus coeruleus. Journal of Comparative Neurology, 155(1), 15–41. 10.1002/cne.901550103

Pierrot-Deseilligny, C., Rivaud, S., Gaymard, B., Müri, R., & Vermersch, A.-I. (1995). Cortical control of saccades. Annals of Neurology, 37(5), 557–567. 10.1002/ana.410370504

Quigley, C., Andersen, S. K., Schulze, L., Grunwald, M., & Müller, M. M. (2010). Feature-selective attention: Evidence for a decline in old age. Neuroscience Letters, 474(1), 5–8. 10.1016/j.neulet.2010.02.053

Quigley, C., & Müller, M. M. (2014). Feature-Selective Attention in Healthy Old Age: A Selective Decline in Selective Attention? Journal of Neuroscience, 34(7), 2471–2476. 10.1523/JNEUROSCI.2718-13.2014

Robinson, O., Vytal, K., Cornwell, B., & Grillon, C. (2013). The impact of anxiety upon cognition: Perspectives from human threat of shock studies. Frontiers in Human Neuroscience, 7. https://www.frontiersin.org/articles/10.3389/fnhum.2013.00203

Sakaki, M., Ueno, T., Ponzio, A., Harley, C. W., & Mather, M. (2019). Emotional arousal amplifies competitions across goal-relevant representation: A neurocomputational framework. Cognition, 187, 108–125. 10.1016/j.cognition.2019.02.011

Sawada, R., & Sato, W. (2015). Emotional attention capture by facial expressions. Scientific Reports, 5(1), 14042. 10.1038/srep14042

Sawada, R., Sato, W., Kochiyama, T., Uono, S., Kubota, Y., Yoshimura, S., & Toichi, M. (2014). Sex differences in the rapid detection of emotional facial expressions. PloS One, 9(4), e94747. 10.1371/journal.pone.0094747

Sawaguchi, T., & Kikuchi, Y. (1998). Noradrenergic effects on activity of prefrontal cortical neurons in behaving monkeys. Advances in Pharmacology (San Diego, Calif.), 42, 759–763. 10.1016/s1054-3589(08)60858-3

Sawaki, R., & Luck, S. J. (2010). Capture versus suppression of attention by salient singletons: Electrophysiological evidence for an automatic attend-to-me signal. Attention, Perception, & Psychophysics, 72(6), 1455–1470. 10.3758/APP.72.6.1455

Schall, J. D. (1995). Neural basis of saccade target selection. Reviews in the Neurosciences, 6(1), 63–85.

Schmitz, A., & Grillon, C. (2012). Assessing fear and anxiety in humans using the threat of predictable and unpredictable aversive events (the NPU-threat test). Nature Protocols, 7(3), 527–532. 10.1038/nprot.2012.001

Schupp, H. T., Stockburger, J., Codispoti, M., Junghöfer, M., Weike, A. I., & Hamm, A. O. (2007). Selective Visual Attention to Emotion. Journal of Neuroscience, 27(5), 1082–1089. 10.1523/JNEUROSCI.3223-06.2007

Schwarz, L. A., Miyamichi, K., Gao, X. J., Beier, K. T., Weissbourd, B., DeLoach, K. E., Ren, J., Ibanes, S., Malenka, R. C., Kremer, E. J., & Luo, L. (2015). Viral-genetic tracing of the input-output organization of a central noradrenaline circuit. Nature, 524(7563), 88–92. 10.1038/nature14600

Smyth, A. C., & Shanks, D. R. (2011). Aging and implicit learning: Explorations in contextual cuing. Psychology and Aging, 26(1), 127–132. 10.1037/a0022014

Sommer, M. A., & Wurtz, R. H. (2002). A Pathway in Primate Brain for Internal Monitoring of Movements. Science, 296(5572), 1480–1482. 10.1126/science.1069590

Sommer, M. A., & Wurtz, R. H. (2006). Influence of the thalamus on spatial visual processing in frontal cortex. Nature, 444(7117), 374–377. 10.1038/nature05279

Stilwell, B. T., & Vecera, S. P. (2022). Testing the underlying processes leading to learned distractor rejection: Learned oculomotor avoidance. Attention, Perception, & Psychophysics, 84(6), 1964–1981. 10.3758/s13414-022-02483-6

Sutherland, M. R., & Mather, M. (2018). Arousal (but not valence) amplifies the impact of salience. Cognition and Emotion, 32(3), 616–622. 10.1080/02699931.2017.1330189

Sutherland, M. R., McQuiggan, D. A., Ryan, J. D., & Mather, M. (2017). Perceptual Salience Does Not Influence Emotional Arousal’s Impairing Effects on Top-Down Attention. Emotion (Washington, D.C.), 17(4), 700–706. 10.1037/emo0000245

Sutton, T. M., & Lutz, C. (2019). Attentional capture for emotional words and images: The importance of valence and arousal. Canadian Journal of Experimental Psychology / Revue Canadienne de Psychologie Expérimentale, 73(1), 47–54. 10.1037/cep0000154

Svärd, J., Fischer, H., & Lundqvist, D. (2014). Adult age-differences in subjective impression of emotional faces are reflected in emotion-related attention and memory tasks. Frontiers in Psychology, 5. 10.3389/fpsyg.2014.00423

Takahashi, J., Shibata, T., Sasaki, M., Kudo, M., Yanezawa, H., Obara, S., Kudo, K., Ito, K., Yamashita, F., & Terayama, Y. (2015). Detection of changes in the locus coeruleus in patients with mild cognitive impairment and Alzheimer’s disease: High-resolution fast spin-echo T1-weighted imaging. Geriatrics & Gerontology International, 15(3), 334–340. 10.1111/ggi.12280

Theeuwes, J., Kramer, A. F., Hahn, S., & Irwin, D. E. (1998). Our Eyes do Not Always Go Where we Want Them to Go: Capture of the Eyes by New Objects. Psychological Science, 9(5), 379– 385. 10.1111/1467-9280.00071

Unsworth, N., & Robison, M. K. (2017). A locus coeruleus-norepinephrine account of individual differences in working memory capacity and attention control. Psychonomic Bulletin & Review, 24(4), 1282–1311. 10.3758/s13423-016-1220-5

van Belle, J., Vink, M., Durston, S., & Zandbelt, B. B. (2014). Common and unique neural networks for proactive and reactive response inhibition revealed by independent component analysis of functional MRI data. NeuroImage, 103, 65–74. 10.1016/j.neuroimage.2014.09.014

Varazzani, C., San-Galli, A., Gilardeau, S., & Bouret, S. (2015). Noradrenaline and Dopamine Neurons in the Reward/Effort Trade-Off: A Direct Electrophysiological Comparison in Behaving Monkeys. Journal of Neuroscience, 35(20), 7866–7877. 10.1523/JNEUROSCI.0454-15.2015

Veríssimo, J., Verhaeghen, P., Goldman, N., Weinstein, M., & Ullman, M. T. (2022). Evidence that ageing yields improvements as well as declines across attention and executive functions. Nature Human Behaviour, 6(1), Article 1. 10.1038/s41562-021-01169-7

Vissers, M. E., van Driel, J., & Slagter, H. A. (2016). Proactive, but Not Reactive, Distractor Filtering Relies on Local Modulation of Alpha Oscillatory Activity. Journal of Cognitive Neuroscience, 28(12), 1964–1979. 10.1162/jocn_a_01017

Vuilleumier, P., Armony, J. L., Driver, J., & Dolan, R. J. (2001). Effects of attention and emotion on face processing in the human brain: An event-related fMRI study. Neuron, 30(3), 829– 841. 10.1016/s0896-6273(01)00328-2

Wagenmakers, E.-J., Love, J., Marsman, M., Jamil, T., Ly, A., Verhagen, J., Selker, R., Gronau, Q. F., Dropmann, D., Boutin, B., Meerhoff, F., Knight, P., Raj, A., van Kesteren, E.-J., van Doorn, J., Šmíra, M., Epskamp, S., Etz, A., Matzke, D., … Morey, R. D. (2018). Bayesian inference for psychology. Part II: Example applications with JASP. Psychonomic Bulletin & Review, 25(1), 58–76. 10.3758/s13423-017-1323-7

Wagenmakers, E.-J., Marsman, M., Jamil, T., Ly, A., Verhagen, J., Love, J., Selker, R., Gronau, Q. F., Šmíra, M., Epskamp, S., Matzke, D., Rouder, J. N., & Morey, R. D. (2018). Bayesian inference for psychology. Part I: Theoretical advantages and practical ramifications. Psychonomic Bulletin & Review, 25(1), 35–57. 10.3758/s13423-017-1343-3

Weinshenker, D. (2018). Long Road to Ruin: Noradrenergic Dysfunction in Neurodegenerative Disease. Trends in Neurosciences, 41(4), 211–223. 10.1016/j.tins.2018.01.010

West, R., & Alain, C. (2000). Age-related decline in inhibitory control contributes to the increased Stroop effect observed in older adults. Psychophysiology, 37(2), 179–189. 10.1111/1469-8986.3720179

Wilcockson, T. D. W., Mardanbegi, D., Xia, B., Taylor, S., Sawyer, P., Gellersen, H. W., Leroi, I., Killick, R., & Crawford, T. J. (2019). Abnormalities of saccadic eye movements in dementia due to Alzheimer’s disease and mild cognitive impairment. Aging, 11(15), 5389–5398. 10.18632/aging.102118

Wilson, R. S., Nag, S., Boyle, P. A., Hizel, L. P., Yu, L., Buchman, A. S., Schneider, J. A., & Bennett, D. A. (2013). Neural reserve, neuronal density in the locus ceruleus, and cognitive decline. Neurology, 80(13), 1202–1208. 10.1212/WNL.0b013e3182897103

Won, B.-Y., & Geng, J. J. (2018). Learned suppression for multiple distractors in visual search. Journal of Experimental Psychology. Human Perception and Performance, 44(7), 1128–1141. 10.1037/xhp0000521

Wöstmann, M., Störmer, V. S., Obleser, J., Addleman, D. A., Andersen, S. K., Gaspelin, N., Geng, J. J., Luck, S. J., Noonan, M. P., Slagter, H. A., & Theeuwes, J. (2022). Ten simple rules to study distractor suppression. Progress in Neurobiology, 213, 102269. 10.1016/j.pneurobio.2022.102269

Yamagishi, S., & Furukawa, S. (2020). Factors Influencing Saccadic Reaction Time: Effect of Task Modality, Stimulus Saliency, Spatial Congruency of Stimuli, and Pupil Size. Frontiers in Human Neuroscience, 14, 571893. 10.3389/fnhum.2020.571893

Ycaza Herrera, A., Wang, J., & Mather, M. (2019). The gist and details of sex differences in cognition and the brain: How parallels in sex differences across domains are shaped by the locus coeruleus and catecholamine systems. Progress in Neurobiology, 176, 120–133. 10.1016/j.pneurobio.2018.05.005

Zanto, T. P., & Gazzaley, A. (2017). Selective attention and inhibitory control in the aging brain. In Cognitive neuroscience of aging: Linking cognitive and cerebral aging, 2nd ed (pp. 207– 234). Oxford University Press. 10.1093/acprof:oso/9780199372935.003.0009

Zeef, E. J., Sonke, C. J., Kok, A., Buiten, M. M., & Kenemans, J. L. (1996). Perceptual factors affecting age-related differences in focused attention: Performance and psychophysiological analyses. Psychophysiology, 33(5), 555–565. 10.1111/j.1469-8986.1996.tb02432.x

Zelinsky, G. J., & Bisley, J. W. (2015). The what, where, and why of priority maps and their interactions with visual working memory. Annals of the New York Academy of Sciences, 1339(1), 154–164. 10.1111/nyas.12606

Zhang, S., Hu, S., Chao, H. H., & Li, C.-S. R. (2016). Resting-State Functional Connectivity of the Locus Coeruleus in Humans: In Comparison with the Ventral Tegmental Area/Substantia Nigra Pars Compacta and the Effects of Age. Cerebral Cortex, 26(8), 3413–3427. 10.1093/cercor/bhv172

Zsidó, A. N. (2023). The effect of emotional arousal on visual attentional performance: A systematic review. Psychological Research. 10.1007/s00426-023-01852-6

Zsido, A. N., Matuz, A., Inhof, O., Darnai, G., Budai, T., Bandi, S., & Csatho, A. (2020). Disentangling the facilitating and hindering effects of threat-related stimuli–A visual search study. British Journal of Psychology, 111(4), 665–682.

